# Selective nigrosome vulnerability in Parkinson’s disease revealed by histology-anchored ultra-high-field quantitative MRI

**DOI:** 10.64898/2026.09.24.754224

**Authors:** Ruiyi Li, Evgeniya Kirilina, Pierre-Louis Bazin, Kjersti Eline Stige, Runa G. Unsgård, Malte Brammerloh, Ingrid Gylterud Kvålsgard, Anne Louise Kristoffersen, Erik Magnus Berntsen, Pål Erik Goa, Philippe Damier, Charalampos Tzoulis, Thanh Pierre Doan

## Abstract

Selective vulnerability is a hallmark of Parkinson’s disease (PD): post-mortem, dopamine-containing cell loss in the substantia nigra first and most severely affects calbindin-poor compartments termed nigrosomes. Testing this in vivo has been limited by circularity, as MRI studies have delineated nigrosomes from the same disease-related contrast they then measure. Here, we warp histology-derived, calbindin-validated probabilistic masks of nigrosomes 1-5 into a cohort-specific multimodal 7 Tesla quantitative MRI template, providing a degeneration-independent anatomical scaffold, in 28 patients with PD and 28 age- and sex-matched controls. In controls, nigrosomes differ from the surrounding nigral matrix in magnetic susceptibility and R1. PD shifts nigrosomal values upward relative to the matrix in susceptibility, R2*, and R1, concentrated in nigrosomes 1, 2, and 4, whereas whole-nigra analysis shows no group difference. Within the matrix, disease effects are largest caudally and emerge dorsally from the ventral tier, and susceptibility alterations in nigrosomes 1 and 4 track contralateral motor symptoms asymmetry. The multimodal pattern suggests that the detectable signal reflects predominantly secondary, iron-associated glial processes rather than neuronal loss itself. These findings recover the post-mortem spatial pattern of nigral vulnerability in vivo and establish a degeneration-independent framework for subregional nigral imaging in PD.

## Introduction

Parkinson’s disease (PD) is among the fastest-growing neurological disorders worldwide^1^ and a leading contributor to neurological disability and economic burden.^2^ Its cardinal motor features of bradykinesia, rigidity, resting tremor and postural instability arise primarily from degeneration of pigmented dopamine-containing neurons in the substantia nigra pars compacta and the subsequent denervation of the nigrostriatal pathway, which disrupts the basal ganglia-thalamocortical circuits that govern movement.^3^

Nigral degeneration in PD is not uniform. From the earliest neuropathological studies, the substantia nigra (SN) has been recognized as regionally heterogeneous, with distinct subregions differing in vulnerability to neurodegeneration. Trétiakoff localized the lesion of PD to the pigmented dopamine-containing neurons of the SN,^4^ and Hassler showed that cell loss falls predominantly on its ventrolateral tier.^5^ Quantitative stereology later confirmed and refined this pattern: pigmented-neuron loss in PD is greatest in the caudal and ventrolateral nigra and follows a gradient distinct from that of normal ageing.^6^ However, each of these frameworks defined vulnerability through the dopamine-containing neurons themselves, which degenerate in PD, leaving no stable anatomical reference against which to map the disease.

This limitation was resolved by a compartmental framework based on calbindin-D28K immunohistochemistry.^7^ By analogy to the striosome-matrix mosaic of the striatum, first defined by histochemical staining and later by calbindin,^8^ Damier and colleagues identified five clusters of pigmented dopamine-containing neurons located in calbindin-poor zones, termed nigrosomes (N1-N5), embedded within a calbindin-rich matrix.^7^ Because this calbindin scaffold is preserved in both health and disease, it provides a degeneration-independent reference onto which cell loss can be mapped directly. Within this framework nigrosomal degeneration in PD follows a stereotyped spatiotemporal sequence, with N1 sustaining the earliest and most severe loss, followed by N2 and N4, while N3 and N5 remain relatively spared.^9^ This nigrosome-centred account of selective vulnerability has since become a cornerstone of our understanding of nigral pathology, but it rests entirely on post-mortem tissue, inferred from a small number of late-stage cases. Whether the same hierarchy describes the in vivo brain has not yet been established.

Testing this in vivo using MRI has, however, proven difficult for reasons as much definitional as technical. Nigrosomes are submillimetre in size, anatomically complex^10^ and embedded in tissue whose MRI contrast mechanisms remain only partly understood.^11^ Most attempts to detect pathological changes in nigrosomes in vivo have departed from their original histological definition in one of two ways. First, the dorsolateral nigral hyperintensity (DNH, swallowtail sign) on susceptibility imaging has been widely treated as an imaging surrogate for nigrosome 1,^12^ even though the two structures do not necessarily coincide anatomically.^11,13,14^ Second, studies that attempt to resolve individual nigrosomes have thus far delineated them from the MRI signal itself, either by visual rating or from voxel-wise patient-versus-control differences.^15–18^ However, defining a nigrosome from the disease-related contrast one intends to measure is circular, operator-dependent and cannot recover nigrosomes that generate little visible signal, precisely the confound that a calbindin-anchored, degeneration-independent definition was designed to avoid. A further constraint is methodological: much of this literature relies on weighted sequences, which are highly scanner- and protocol-dependent and not directly comparable across sites or time points.^17,18^ By contrast, quantitative MRI (qMRI), such as quantitative susceptibility mapping (QSM), the effective transverse relaxation rate (R2*), and the longitudinal relaxation rate (R1), yields reproducible, physically defined measures whose differential sensitivity to iron and myelin permits biological interpretation of the underlying tissue rather than mere signal contrast.^19,20^

Here we address both the definitional and the methodological constraint. We warp histology-derived, calbindin-validated probabilistic masks of nigrosomes N1-N5^21^ into a cohort-specific multimodal ultra-high-resolution qMRI template at 7 Tesla (7 T), establishing in vivo a degeneration-independent anatomical scaffold onto which iron- and myelin-sensitive measures from patients and matched controls are mapped. This recovers the post-mortem vulnerability hierarchy in living patients, localises it to N1, N2 and N4, links nigrosomal susceptibility asymmetry to clinically observed contralateral motor asymmetry, and constrains what the nigral qMRI signal at this disease stage is likely to reflect.

## Materials and methods

### Participants

A total of 58 participants, 29 subjects with PD and 29 age- and sex-matched healthy controls, were initially included in this study. Twenty-nine patients and nine healthy controls (HCs) were drawn from the STRAT-PARK cohort^22^, and a further 20 HCs, selected to age- and sex match the total HC distribution to the PD distribution, were drawn from an ongoing quality assurance study at the same site. All 58 subjects, regardless of source cohort, were recruited at St. Olav’s University Hospital, Trondheim, Norway, and scanned on the same scanner, using identical protocols (mGRE, MP2RAGE, and B1+ mapping) and postprocessing pipelines (described below).

Patients were recruited by referral from the outpatient clinic. Inclusion criteria for patients included a diagnosis of clinically established or clinically probable PD according to the Movement Disorders Society (MDS) diagnostic criteria for PD^23^, ability to consent and participate in the study, and a DAT scan or ¹⁸F-FDOPA scan showing a loss of dopamine nerve endings in the striatum. Motor functions were assessed using the MDS Unified Parkinson’s Disease Rating Scale (MDS-UPDRS) part III in the “ON” dopaminergic state. Hoehn and Yahr staging was likewise performed in the “ON” state. Clinical parameters such as age, sex, disease duration from motor symptom onset, time since last dose of levodopa, and levodopa-equivalent daily dose (LEDD) were all recorded. Controls from STRAT-PARK were primarily spouses or close relatives of patients, without neurodegenerative disease. For controls from the quality assurance cohort, self-reported central nervous system disease or conditions affecting brain iron concentration, including neurodegenerative, neuroinflammatory, genetic, and systemic disorders, were exclusion criteria. The sample size was determined by the number of eligible participants with usable imaging and clinical data available from the source cohorts; therefore, no a priori sample-size calculation was performed.

As a sensitivity analysis, the differences in mean SN signal between the two independent control cohorts were tested with OLS regression with cohort as the predictor of interest. Moreover, four controls within the quality assurance cohort with incidental radiological signs of small vessel disease, defined as white-matter changes of Fazekas grade ≥2 below 75 years of age or ≥3 at 75 years or older, and/or central microhaemorrhages or lacunar infarcts^24^ (all four were older than 75 years with Fazekas grade 3), were tested against the rest of the quality assurance cohort using the same procedure, but with group as the predictor of interest.

The study is in full accordance with the Declaration of Helsinki and has been approved by the Norwegian ethics committee (REK 74985 and REK 108066). All included participants have provided written informed consent.

### Motor lateralization scoring

Motor lateralization was quantified from item-level MDS-UPDRS III scores. Right-sided scores were computed as the sum of right-sided bradykinesia items (finger tapping, hand movements, pronation-supination, leg agility and toe tapping), right-sided tremor items (postural tremor, kinetic tremor and two resting tremor items for hand and leg) and right-sided rigidity items (arm and leg). Left-sided scores were computed identically from the corresponding left-sided items. Midline symptoms such as axial items (speech, facial expression, arising from chair, posture, gait, postural instability and freezing), neck rigidity and midline tremor items (lip/jaw resting tremor, constancy of resting tremor) were summed separately and used to compute a total MDS-UPDRS III score (the sum of the lateralized and midline item scores), but were not included in the lateralized scores.

### MRI acquisition

All subjects were scanned using the same 7 T Siemens MAGNETOM Terra scanner and 1Tx/32Rx head coil. The imaging protocol included a 3D multi-gradient recalled echo (mGRE), a 3D Magnetization Prepared 2 Rapid Acquisition Gradient Echoes (MP2RAGE) sequence^25^, and a B1+ flip-angle mapping sequence, which were further processed for the calculation of QSM, R2*, and R1. Three-dimensional mGRE with four echoes in the transverse plane and a voxel size of 0.75 mm isotropic was acquired using the ASPIRE framework^26^: repetition time = 31 ms, echo times = 2.54/7.22/14.44/23.23 ms, bandwidth = 300/300/200/100 Hz/px, flip angle = 12°, matrix = 304 × 228 × 224, GRAPPA acceleration factor R = 3. Three-dimensional MP2RAGE in the sagittal plane with a voxel size of 0.75 mm isotropic was acquired with the following parameters: echo time = 1.99 ms, inversion TR/excitation TR = 4300 ms/7.2 ms, inversion time 1/inversion time 2 = 840 ms/2370 ms, bandwidth = 250 Hz/px, flip angle 1/flip angle 2 = 5°/6°, matrix = 224 × 300 × 320, excitations = 180, GRAPPA acceleration factor R = 3. Two-dimensional B1+ flip-angle maps in the transverse plane were acquired using a pre-saturated TurboFLASH B1+ mapping sequence (tfl_b1map)^27^ with the following parameters: repetition time = 4000 ms, echo time = 1.72 ms, flip angle = 90°, slice thickness/spacing = 4/8 mm, matrix = 64 × 64, bandwidth = 490 Hz/px, GRAPPA acceleration factor R = 2.

### Quantitative MRI processing

Calculation of qMRI maps was performed using an in-house Python script based on the Nighres library version 1.5.2.^28^ Prior to quantitative fitting, the multi-echo magnitude and phase images were denoised using local complex PCA (neighbourhood size 4, standard deviation cutoff 1.05) with integrated phase unwrapping and rescaling.^29^ Brain masking was performed on the N4ITK bias-field-corrected MP2RAGE INV2 image using intensity-based skull stripping.^30^ Rigid co-registration parameters between the first-echo mGRE magnitude and the MP2RAGE INV2 image were then estimated and used to bring MP2RAGE space images into mGRE space. The resampled brain mask was used for all subsequent masking.

QSM was reconstructed from echoes 2-4 using a Python implementation of the TGV QSM algorithm.^31^ Phase images were rescaled from scanner units to radians, and the brain mask was additionally eroded by five voxels to minimize boundary artefacts at the brain-air interface. TGV QSM was run for 1000 iterations with regularization parameters α = 0.0015 and 0.0005. QSM was reconstructed independently from each echo, and the final map was obtained as the voxelwise median across echoes, clipped to [−0.3, 0.5] ppm. R2* maps were computed from the four denoised magnitude echoes by log-linear least-squares fitting of a monoexponential decay model, with values clipped to a maximum of 200 s⁻¹ after brain masking. R1 maps were derived from the MP2RAGE uniform image with a lookup table based on imaging parameters. B1 maps were resampled onto the MP2RAGE UNI grid using the acquisition header geometry and used to correct R1. R1 was then resampled into mGRE space with the inverse of the MP2RAGE mGRE coregistration and clipped to a maximum of 3 s⁻¹ after brain masking. The resulting qMRI maps were all visually inspected for reconstruction quality and artefacts.

### Multimodal template construction

A cohort-specific multimodal template was created with the Advanced Normalization Tools (ANTs) environment version 2.6.3 using iterative template creation with joint cross-correlation over all three modalities (Fig. 1C).^32^ To ensure a good initial alignment, all 56 subjects (28 patients and 28 HCs) included in the final cohort were first prealigned to a common reference by rigid and affine registration of the QSM volumes, driven by a mutual information metric (32 bins, regular sampling, fraction 0.25) and optimized across four resolution levels (shrink factors 8×4×2×1, smoothing 3×2×1×0 vox, convergence threshold 1 × 10⁻⁶ over 1000×500×250×100 iterations). The resulting transforms were propagated to the R1 and R2* volumes of each subject, and an initial template was computed per modality as the voxelwise mean of the prealigned images. Iterative multivariate template refinement was initialized from these averages, with antsMultivariateTemplateConstruction2.sh over four iterations, adapting the multimodal weighting strategy of a prior study.^33^ At each iteration, each subject was registered to the current template using rigid and affine initialization (mutual information; 1000×500×250×0 iterations, shrink factors 6×4×2×1, smoothing 4×2×1×0 vox, convergence threshold 1 × 10⁻⁶) followed by symmetric diffeomorphic normalization (SyN[0.10, 3, 0]) driven by cross-correlation (neighbourhood radius 4) jointly across all three modalities (weights QSM 1.0, R2* 1.0, R1 0.7, one deformation field per subject). The SyN stage used four resolution levels (shrink factors 6×4×2×1, smoothing 3×2×1×0 vox, maximum 100×70×50×20 iterations, convergence threshold 1 × 10⁻⁹). At each iteration, the template was updated toward the population mean with a shape-update gradient step of 0.08. Following template construction, a final shape update with a gradient step of 0.25 was applied to each modality template. The resulting templates (Supplementary Fig. 1) and subjects in template space were all visually inspected for misalignments and artefacts.

**Fig. 1:**
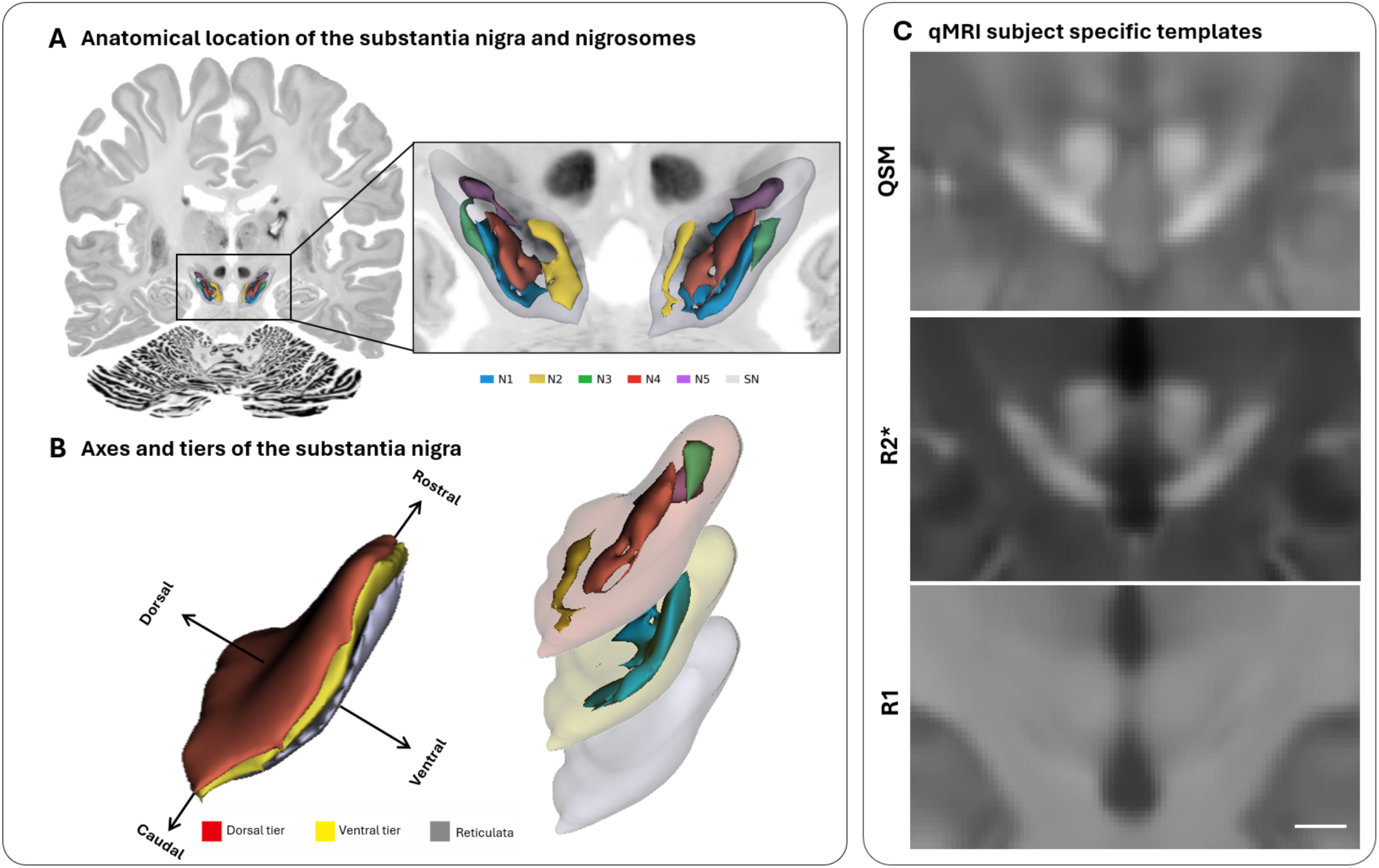
Anatomy of the substantia nigra and multimodal cohort-specific qMRI templates. **(A)** Whole-brain coronal section showing the anatomical location of the SN and nigrosomes 1-5 (N1-N5) co-registered to the BigBrain single-subject histological volume in MNI ICBM 2009b asymmetric space at 300 µm.^34^ Nigrosomes 1-5 (N1-N5) are colour-coded according to the legend and displayed within a translucent SN volume. **(B)** Three-dimensional representation of layers of the SN along the dorsoventral axis. The upper layer (dorsal tier) and intermediate layer (ventral tier) contain dopamine-rich but calbindin-poor nigrosomes, surrounded by calbindin-rich neuropil, and are commonly referred to as the pars compacta. Nigrosomes 2-5 are located within the dorsal tier, whereas nigrosome 1 is located within the ventral tier. The outermost ventrolateral layer (grey) contains no nigrosomal tissue and is commonly referred to as pars reticulata. It consists predominantly of GABAergic neurons surrounded by calbindin-positive neuropil. Note, however, that pars compacta and pars reticulata are defined by cell type (dopaminergic and GABAergic neurons, respectively) rather than by an anatomical boundary. The layers do therefore not delineate these divisions, but rather approximate that the dopaminergic neurons (pars compacta) are largely confined to the dorsal and ventral tiers, and the GABAergic neurons (pars reticulata) to the outermost ventrolateral layer. The rostrocaudal and dorsoventral axes are indicated; the mediolateral axis is orthogonal to both. **(C)** Coronal section of the cohort-specific quantitative susceptibility mapping (QSM), effective transverse relaxation rate (R2*) and longitudinal relaxation rate (R1) templates at the level of the SN. All contrast profiles are presented at the same spatial scale. Scale bar, 7.5 mm.

### Multimodal focused nigrosome atlas registration

Histology-derived probability masks of nigrosomes 1-5 and whole SN in the AHEAD atlas template space were warped into the cohort-specific template space using a Nighres-based focused multi-contrast registration script provided in the publication.^21^ QSM, R2* and R1 templates were co-registered to their corresponding AHEAD templates in a two-stage multichannel fit, with each modality contributing equally to a mutual information cost. A global alignment step using rigid, affine, and SyN transformations was used to move the two template spaces into the same space, followed by a refined SyN stage focused on a 3 mm neighbourhood region around the SN probability mask in AHEAD, limiting the refined nonlinear deformation to the midbrain rather than fitting a whole-brain warp. The SyN stage used a coarse-to-fine schedule (40 coarse, 50 medium, 40 fine iterations) with high regularization in Stage 1 and low regularization in Stage 2, and nearest-neighbour resampling during optimization. Mask overlap and size diagnostics can be found in the Supplementary material (Supplementary Fig. 2 and Supplementary Table 1).

### Region of interest value extraction

Quantitative map values were extracted from each ROI as probability-weighted means. Where appropriate, lateralized ROIs were created by midline split, and values were extracted separately from left and right masks. A probabilistic pooled nigrosome mask was derived as the voxelwise maximum across the five individual nigrosome probability maps, max (N1, N2, …, N5). The matrix mask was calculated as whole-SN probability minus the pooled nigrosome mask, with negative values clipped to zero, consistent with the original publication in which the matrix is defined as the calbindin-rich territory of the SN surrounding the calbindin-poor nigrosomes.^7^ The matrix therefore contains five tissue components: the dopamine-containing neurons lying outside the nigrosomes, the GABAergic neurons of pars reticulata, striatonigral fibres traversing the nigra, glial cells, and, predominantly, calbindin-positive neuropil. Of these, only the dopamine-containing neurons and glial cells are expected to change with disease, so the matrix region of interest is compositionally heterogeneous by its nature. QSM was CSF-referenced to remove global offsets by subtracting the mean susceptibility within a bilateral lateral ventricle mask acquired from the probabilistic Harvard–Oxford Subcortical Structural Atlas^35–38^ distributed within FSL, thresholded to >0.95 and warped to template space. R2* and R1 were used in their native quantitative units.

### Statistical analysis

#### Statistical framework

Statistical analyses were performed in Python version 3.9.23 except for linear mixed model (LMM) fitting, which was done in R version 4.5.2. All analyses were controlled for age and sex by adding them as covariates, and effect sizes were covariate-adjusted throughout and reported both in native units and standardized (numerator-adjusted Hedges’ *g* and *β*). Ninety-five per cent confidence intervals were reported as stratified bootstraps with 5000 resamples (within-group for case–control analyses). Inference on coefficients of interest was two-sided with Freedman-Lane permutation (n = 10 000) of the reduced-model residuals for ordinary least squares (OLS) regression, and with parametric-bootstrap likelihood-ratio test (n = 10 000) for LMM. PD vs HC analysis had 52 degrees of freedom, while PD-only analysis had 24. False discovery rate (FDR) was controlled by the Benjamini-Hochberg procedure on these inferences, within predefined families for each analysis below. FDR-corrected *P* values denoted as *q* were reported throughout, with a significance level of *q* ≤ 0.05. Statistical significance was denoted as n.s. (not significant), +0.05 ≤ *q* < 0.10, \**q* ≤ 0.05, \*\**q* ≤ 0.01, \*\*\**q* ≤ 0.001. Full detailed statistical descriptions were reported in the Supplementary Material.

#### Pooled nigrosomes versus matrix contrast

For each modality and group, the within-subject pooled nigrosome-to-matrix contrast was tested against zero by a one-sample *t*-test on the paired contrast. The PD vs HC difference in pooled nigrosome-to-matrix gap was tested separately across each modality by OLS of the contrast on group. The PD vs HC contrast in pooled nigrosomes and in the matrix was also assessed by the same OLS framework. FDR was applied within four disjoint families: the six within-group gap vs zero tests across modality × group cells, the three PD vs HC modality-level contrasts, and the three nigrosome-signal and three matrix-signal tests, corrected separately within each region.

#### Per-nigrosome group comparisons

Bilateral per-nigrosome values were tested for PD vs HC differences in OLS regressions with group as the predictor of interest. Modality-standard and matrix-subtracted schemes were applied in parallel to analyse bulk effect and the effect after internal reference to the matrix to remove between-subject noise as a sensitivity analysis. FDR was applied across the five nigrosomes within each modality × scheme. Outside the FDR family, whole SN and matrix-referenced SN were reported as a context ROI for modality-standard referencing and matrix referencing, respectively. A threshold-sweep was performed with mask thresholds of 0.2, 0.3, 0.5 and with the upper 25th-percentile of mask probability (labelled core_p75 in Supplementary Fig. 3).

#### Spatial gradients within the matrix

To test the spatial gradient of matrix signal between groups along the three intrinsic axes, an LMM was fit per modality and axis. The three intrinsic SN axes were derived per hemisphere from the SN-mask voxels with the mask thresholded at a probability of 0.2 to restrict the geometry to voxels confidently within the SN. The shortest axis, assigned as the dorsoventral axis, was derived as a Laplace-equation depth field to accommodate curvature of the SN sheet. The two long axes were derived by principal component analysis (PCA) and assigned to the rostrocaudal and mediolateral axes by eigenvalue ordering. Axis directions were fixed by visual assessment. Each axis was range-normalized to [0, 1] within each hemisphere and divided into 10 quantile-based bins.

Subject-level probability-weighted means were extracted per bin within the matrix and modelled with an LMM as

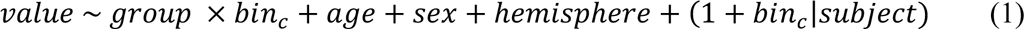

where bin_c_ is centered at the midpoint (5.5) and group × bin_c_ is the coefficient of interest.

Both hemispheres were entered jointly, with hemisphere as a fixed effect and a by-subject random intercept and random slope for bin_c_ linking each participant’s two hemispheres. Models were fit by restricted maximum likelihood (REML; lme4), with reduction to a random intercept when the maximal model was singular or failed to converge. The group × bin_c_ interaction was assessed by a parametric-bootstrap likelihood-ratio test (pbkrtest, 10 000 simulations) comparing maximum-likelihood refits of the full and reduced models, with Satterthwaite approximation (lmerTest) on the REML fit as a sanity check. BH-FDR was applied within each modality across the three axis-interaction tests.

#### Associations with disease duration

The qMRI contrast was regressed on disease duration within PD to track per-nigrosome progression with disease duration. FDR was applied across five nigrosomes within each modality. To display the progression from controls across disease duration, bilateral nigrosome values for all patients in both standard and matrix-referenced schemes were normalized to an HC baseline, expressed as a *z*-score relative to the HC residual standard deviation. Patients were then divided into balanced tertiles based on disease duration, and the mean deviation from HC was reported for each tertile.

#### Imaging and motor asymmetry

Within patients only, OLS regressions tested whether per-modality imaging asymmetry (left − right ROI value) correlated with contralateral motor asymmetry (right − left MDS-UPDRS III lateralized score) per nigrosome. FDR was applied across the five nigrosomes within each modality. Leave-one-out jackknife stability checks and clinical covariate sensitivity analyses (disease duration, time since levodopa, levodopa daily dose) were performed per nigrosome (Supplementary Table 14 and Supplementary Fig. 8).

## Results

### Cohort characteristics

Of the 29 patients, one patient was excluded after visual quality control due to excessive artefacts after qMRI reconstruction. The age- and sex-matched HC was thereafter excluded to balance the cohorts. The resulting PD cohort comprised of 28 subjects, 13 females and 15 males, with a mean age of 66.8 years. The resulting HC group included 28 age- and sex-matched controls, 13 females and 15 males, with a mean age of 66.0 years. The PD and HC groups did not differ in age (Welch’s *t*-test, *P* = 0.731) or in sex (χ² test, *P* = 1.000). The patients ranged in MDS-UPDRS III from 9 to 51 (mean = 25.3), disease duration from 1.3 to 28.2 years (mean = 8.1) and Hoehn and Yahr stage from 1 to 3 (mean = 2.0). Clinical motor dominance (defined as a nonzero difference between right- and left-sided MDS-UPDRS III scores) was left-sided in 14 patients, right-sided in 11, and symmetric in three. Twenty-seven of the 28 patients were on levodopa at the time of assessment, with a mean LEDD of 652 mg/day (range = 300-1780 mg/day) and a median time since last levodopa of 124 min (range = 15-225 min). LEDD was missing for two patients and time since levodopa for one. Complete aggregated statistics can be found in Supplementary Tables 2 and 3.

No significant differences were found in mean SN signal between the two independent control cohorts in any modality, and no differences in mean SN signal were found between the controls with small vessel disease and the rest of the quality assurance cohort in any modality (Supplementary Tables 4 and 5).

### Nigrosome-matrix contrast is amplified in Parkinson’s disease

Pooled nigrosome signal differed from matrix signal within subjects in all three modalities (Fig. 2A and 2B; Supplementary Table 6). In PD, within subject pooled nigrosome signal exceeded the matrix signal by a mean of 0.0336 ppm for QSM (*q* < 0.001) and 3.32 s⁻¹ for R2* (*q* < 0.001), whereas R1 was 0.0237 s⁻¹ lower in the pooled nigrosomes than in the matrix (*q* < 0.001). Controls showed the same pattern for QSM (+0.0234 ppm, *q* < 0.001) and R1 (−0.0318 s⁻¹, *q* < 0.001). The R2* difference in controls was, however, not significant after FDR correction (+0.60 s⁻¹, *q* = 0.286).

**Fig. 2:**
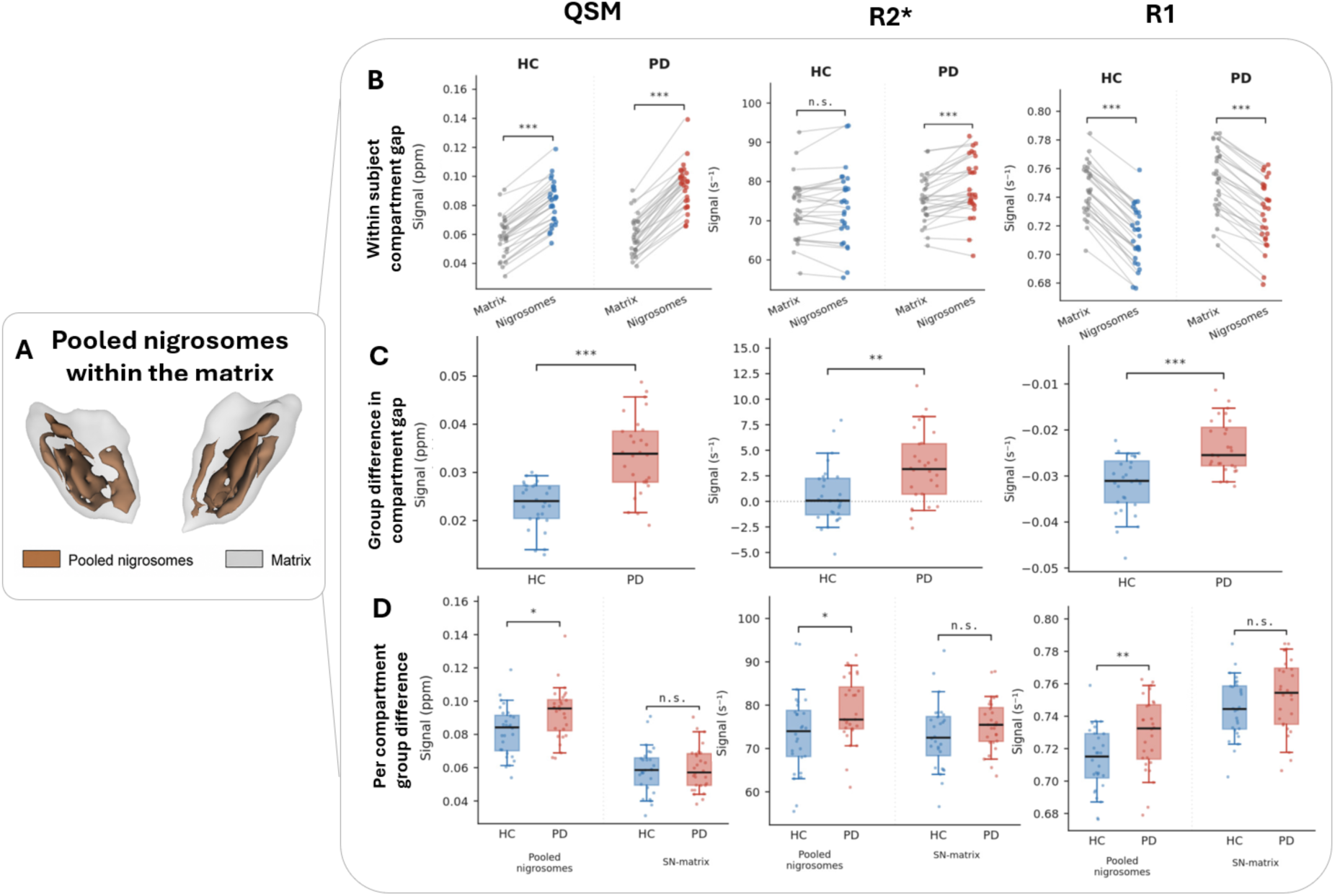
Compartment-specific qMRI differences between pooled nigrosomes and matrix tissue in Parkinson’s disease. **(A)** Three-dimensional rendering of the pooled nigrosomes (brown) within the surrounding matrix (gray). The matrix was defined as the SN after subtraction of the pooled nigrosome masks, consistent with the original definition of the matrix as the calbindin rich territory of the SN surrounding the calbindin-poor nigrosomes.^7^ **(B)** Paired within-subject comparisons showed higher QSM and R2* signals, but lower R1 signals, in the pooled nigrosomes compared to the matrix. This compartmental pattern was present in both HCs and participants with PD. Lines connect measurements from the same participant. **(C)** The nigrosome-to-matrix signal gap differed significantly between groups. Compared with healthy controls, participants with PD showed a larger positive difference for QSM and R2* and a less negative difference for R1. **(D)** Compartment-specific group comparisons showed higher pooled nigrosome signals in PD across all three modalities, whereas no significant group differences were detected in the matrix. Group comparisons are shown with box and whiskers plots displaying the median and interquartile range with the whiskers extending to 1.5 × interquartile range. Red indicates the PD group, and blue indicates the HC group. Each participant is displayed as a datapoint. Statistical significance is denoted as n.s. (not significant), +0.05 ≤ *q* < 0.10, \**q* ≤ 0.05, \*\**q* ≤ 0.01, \*\*\**q* ≤ 0.001

Moreover, nigrosome-matrix contrast differed between PD and controls in all three modalities after FDR correction (Fig. 2C; Supplementary Table 7). PD showed a 0.0101 ppm larger QSM gap (*g* = 1.52, *q* < 0.001) and a 2.70 s⁻¹ larger R2* gap (*g* = 0.83, *q* = 0.003). For R1, the difference was 0.0079 s⁻¹ less negative in PD than in controls (*g* = 1.30, *q* < 0.001).

These changes were driven predominantly by nigrosome signal, rather than matrix signal (Fig. 2D). Compared with controls, PD showed higher pooled nigrosome signal by 0.0108 ppm for QSM (*g* = 0.69, *q* = 0.017), 5.22 s⁻¹ for R2* (*g* = 0.60, *q* = 0.028) and 0.0157 s⁻¹ for R1 (*g* = 0.72, *q* = 0.008). We found no significant group differences in the matrix (Supplementary Table 8).

### Group differences localize to N1, N2, and N4

When examining individual nigrosomes under standard referencing, N1, N2, and N4 showed higher signal in PD than in controls, whereas N3, N5, and the whole SN reference showed no significant group difference (Fig. 3A, C, E). In N1, PD–control differences were found in all three modalities, with effects of +0.0101 ppm for QSM (*g* = 0.69, *q* = 0.028), +5.66 s⁻¹ for R2* (*g* = 0.66, *q* = 0.045), and +0.0179 s⁻¹ for R1 (*g* = 0.82, *q* = 0.003). N4 also differed in all modalities: +0.0186 ppm for QSM (*g* = 1.09, *q* = 0.001), +11.06 s⁻¹ for R2* (*g* = 1.07, *q* = 0.002) and +0.0197 s⁻¹ for R1 (*g* = 0.89, *q* = 0.003). N2 differed only in QSM (+0.0136 ppm; *g* = 0.61, *q* = 0.042). The whole-substantia-nigra reference, N3 and N5, showed no significant group differences in any modality (Supplementary Table 9).

**Fig. 3:**
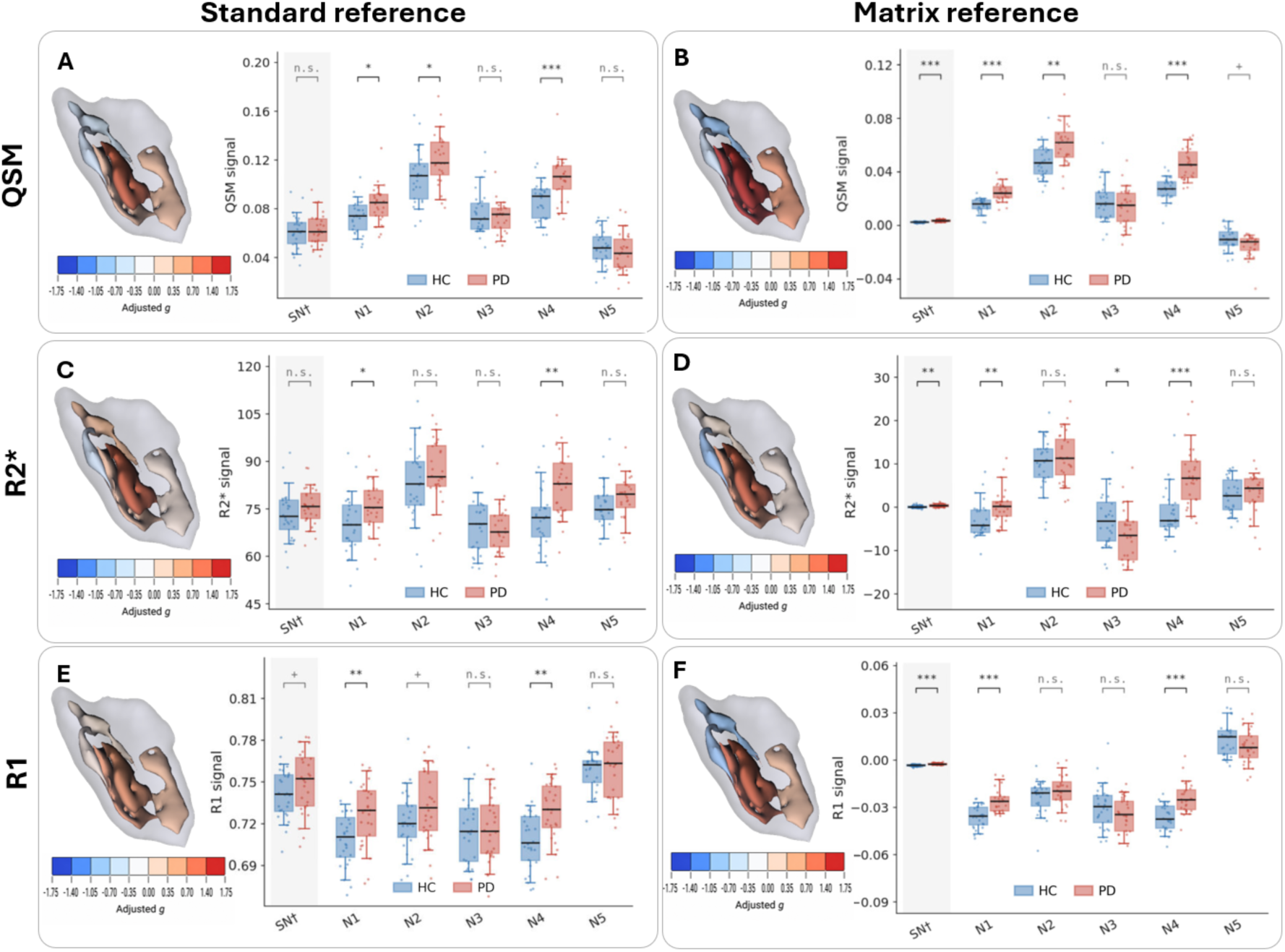
Per-nigrosome differences between Parkinson’s disease and healthy controls under standard and matrix referencing. **(A, C, E)** With standard referencing, participants with PD showed higher signals in N1 and N4 across QSM, R2*, and R1, with an additional increase in N2 for QSM. **(B, D, F)** After referencing to the matrix, the same core pattern of N1, N2, and N4 is preserved, with additional effects in the opposite direction in N3 (R2*) and N5 (QSM). Three-dimensional renderings show the age- and sex adjusted Hedges’ *g* for each nigrosome, with red indicating higher values and blue indicating lower values in PD relative to HC. Box-and-whisker plots displaying the median and interquartile range, with the whiskers extending to 1.5 × interquartile range, show the corresponding participant-level distributions. Only the left SN is displayed, although effects were bilateral. †Reference region. Standard referencing denotes native quantitative units (CSF-referenced QSM; R2* and R1 unreferenced); matrix referencing denotes subtraction of each participant’s mean matrix signal from each regional value. SN denotes the whole substantia nigra. Statistical significance is denoted as n.s. (not significant), +0.05 ≤ *q* < 0.10, \**q* ≤ 0.05, \*\**q* ≤ 0.01, \*\*\**q* ≤ 0.001.

The matrix subtraction sensitivity analysis amplified the same pattern (Fig. 3B, D, F). N1 showed PD–control differences of +0.0094 ppm for QSM (*g* = 1.56, *q* < 0.001), +3.15 s⁻¹ for R2* (*g* = 0.77, *q* = 0.009) and +0.0101 s⁻¹ for R1 (*g* = 1.35, *q* < 0.001). N4 showed differences of +0.0179 ppm for QSM (*g* = 1.72, *q* < 0.001), +8.55 s⁻¹ for R2* (*g* = 1.37, *q* < 0.001) and +0.0119 s⁻¹ for R1 (*g* = 1.28, *q* < 0.001). N2 showed effects in QSM of +0.0129 ppm (*g* = 0.90, *q* = 0.001).

After internal referencing to the matrix, the whole substantia nigra also differed between groups; because such referencing compresses between-subject variance by design, this is reported in the Supplementary material rather than as an independent effect (Supplementary Table 10). Two opposite-direction effects also reached significance: N3 in R2* (−4.09 s⁻¹, g = −0.61, q = 0.037) and N5 in QSM (−0.0047 ppm, g = −0.55, q = 0.050), both likely partial-volume effects. The results were stable across probability thresholds (Supplementary Fig. 3).

### Group differences within the matrix increase towards the caudal and dorsal ends

The matrix showed rostrocaudal and dorsoventral group gradients (Fig. 4A), with larger PD– control differences towards the caudal and dorsal ends (Fig. 4B-D). Along the rostrocaudal axis, the PD–control difference decreased from caudal to rostral for both QSM (*β* = −0.00273, *q* < 0.001) and R1 (*β* = −0.00189, *q* < 0.001). R2* showed a similar caudal pattern, but the rostrocaudal slope was not significant (*β* = −0.288, *q* = 0.389) (Supplementary Table 11 and Supplementary Fig. 4). Along the dorsoventral axis, the PD to HC difference in R2* increased towards the dorsal end (*β* = +0.787, *q* < 0.001). The difference was close to zero in the most ventral bins and emerged from the level corresponding to the ventral tier upwards, consistent with the layer organization of Fig. 1B. No modality showed a significant mediolateral interaction (Supplementary Table 11 and Supplementary Fig. 4). The group profiles overlapped closely for QSM except along the rostrocaudal axis, whereas R2* and R1 were generally higher in PD, consistent with the direction of matrix group differences in the pooled analysis. All LMMs converged for both intercept and slope, except for dorsoventral for QSM and R2* (Supplementary Table 11).

**Fig. 4:**
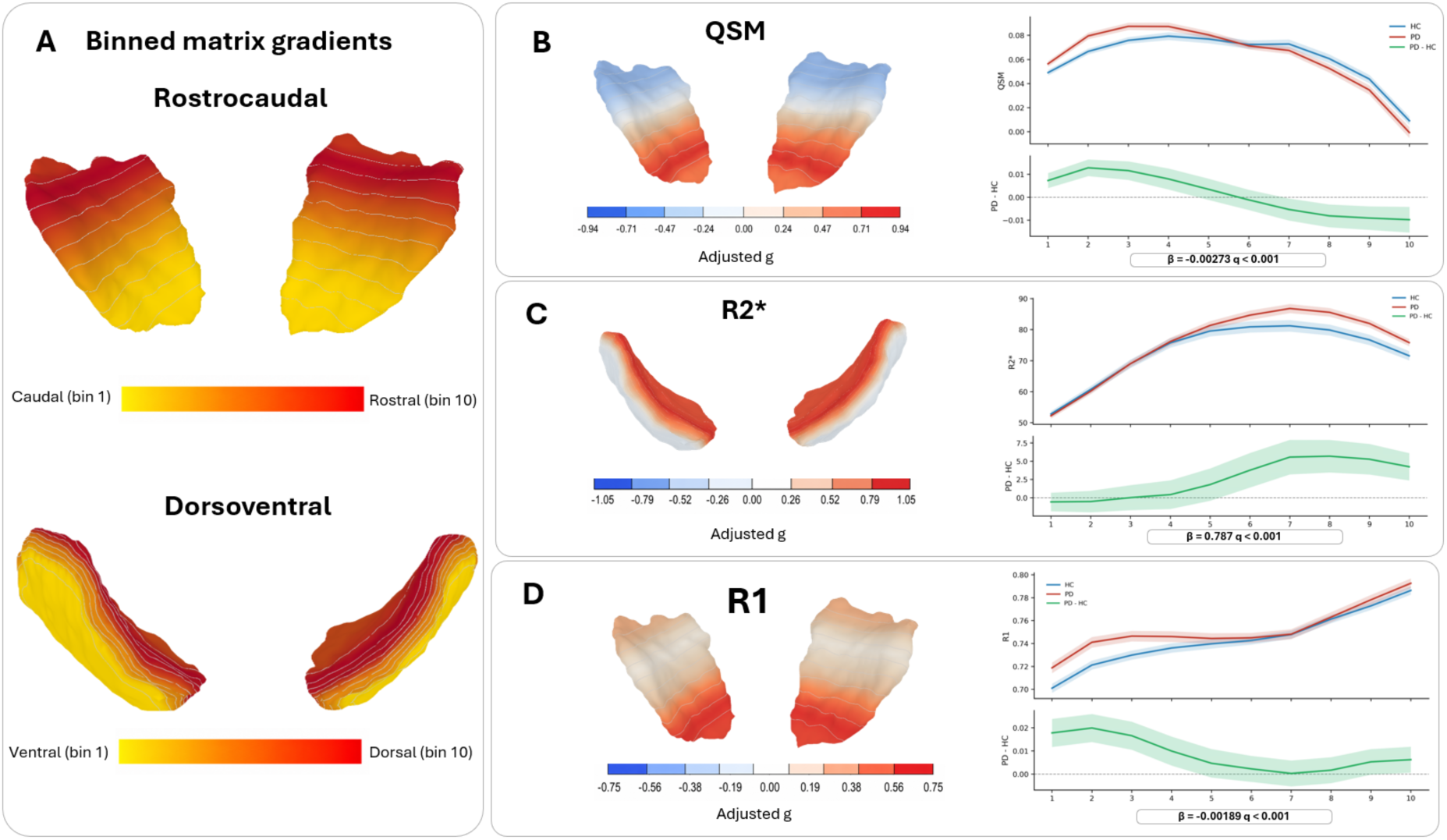
Group × axis interactions in the matrix along rostrocaudal and dorsoventral gradients. (**A**) The rostrocaudal and dorsoventral gradients and bins are displayed on a three-dimensional mesh of the matrix. Bins run from caudal and ventral (bin 1, yellow) to rostral and dorsal (bin 10, red) (**B-D**) Interactions are found in the rostrocaudal axis for both QSM and R1, and a dorsoventral interaction is found in R2*. The largest PD to HC increases were at the caudal end; along the dorsoventral axis, the difference emerged from the level corresponding to the ventral tier and increased dorsally. Effect sizes per bin are displayed as adjusted Hedges’ g on surface renderings of the matrix, with red showing positive effect sizes, and blue showing negative effect sizes. Line plots show group means and 95% confidence bands across bins for PD (red) and HC (blue) and the PD−HC difference (green). Interaction coefficients from the linear mixed models and FDR corrected *q* values are given beneath each plot.

### Associations with disease duration

Across modalities and referencing schemes, contrast in N1, N2 and N4 tended to increase with longer disease duration, with the largest estimate in N4 (Supplementary Tables 12 and 13). However, none of the duration associations survived FDR correction, so these patterns should be interpreted descriptively. The tertile visualization showed a corresponding pattern: N1, N2 and N4 were above the healthy-control baseline, with deviations increasing with disease duration in all modalities, whereas N3 and N5 were below the baseline (Fig. 5A-F and Supplementary Fig. 5; per participant values in Supplementary Fig. 6).

**Fig. 5:**
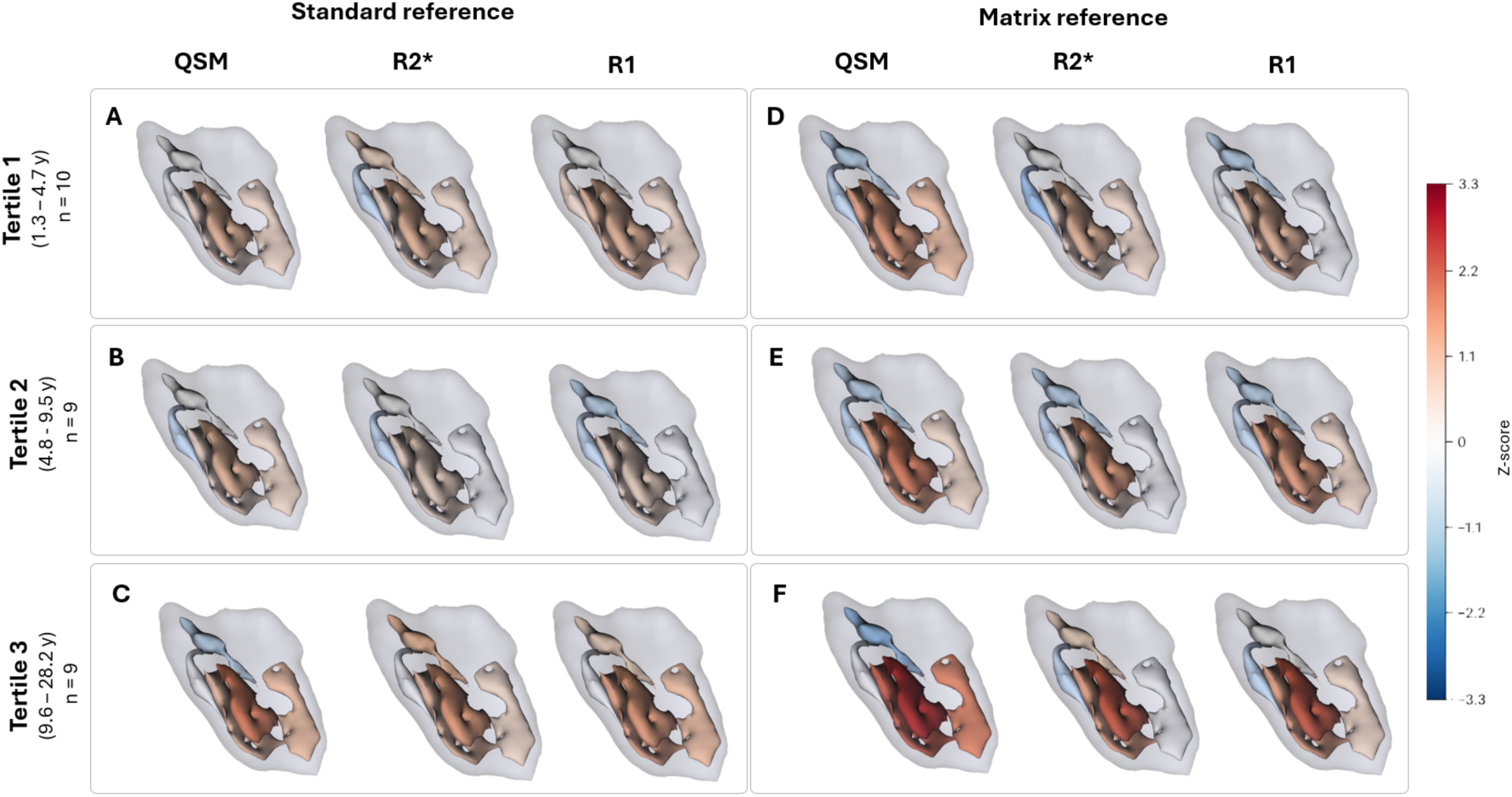
Per nigrosome deviation from the healthy control mean across disease duration tertiles. **(A-C)** Deviation from the healthy control mean, expressed as an age- and sex-adjusted z-score per nigrosome, was most pronounced in N1, N2, and N4 across all modalities and tertiles under modality standard reference and tended to increase with disease duration. No duration slope did however survive FDR correction, so the observation is only descriptive. Because z-scores scale each deviation by the healthy-control variability of that region, they index deviation relative to normal variation rather than absolute effect size. (**D-F**) The same pattern is preserved after referencing to the matrix. Three-dimensional renderings show the age- and sex adjusted z score for each nigrosome and tertile, with red indicating positive values and blue indicating negative values. Tertile boundaries and group sizes are given at the left of each row.

### Susceptibility asymmetry in N1 and N4 correlates with contralateral motor asymmetry

Imaging asymmetry tracked clinical motor asymmetry in a spatially specific pattern that mirrored the group-level topography, with N1 and N4 showing significant correlations in QSM (Fig. 6). N1 showed the strongest association (+451.8 MDS-UPDRS III points per ppm, equivalent to 4.5 points per 0.01 ppm of susceptibility asymmetry; standardized *β* = 0.645, *q* = 0.002), followed by N4 (+253.0 points per ppm, or 2.5 points per 0.01 ppm; standardized *β* = 0.538, *q* = 0.010). N2, N3, and N5 showed no significant associations.

**Fig. 6:**
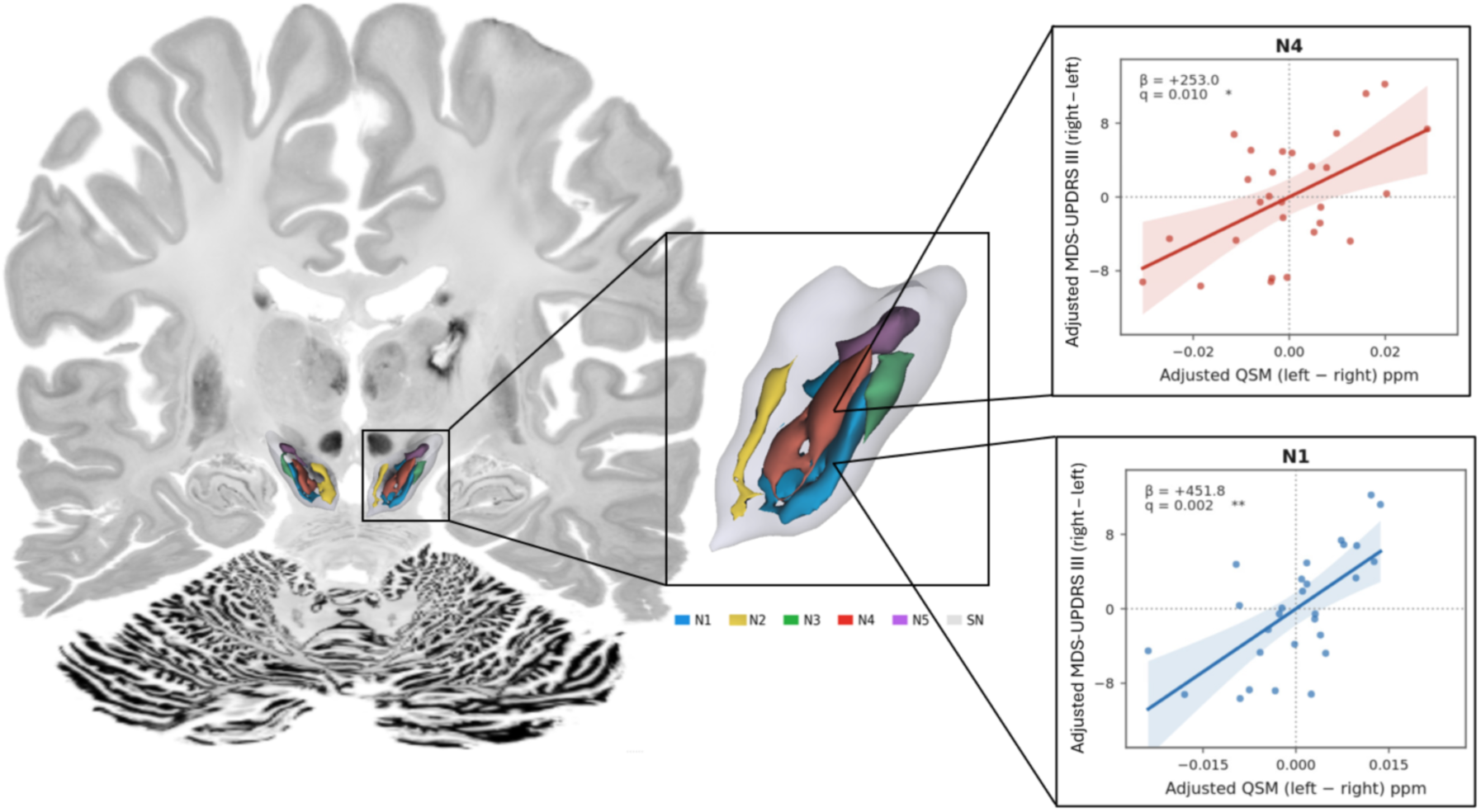
Correlation between asymmetry in contralateral susceptibility and MDS-UPDRS III. Correlations between asymmetry in contralateral QSM signal and MDS-UPDRS III were found in both N1 (blue) and N4 (red), with the largest slope found in N1. Regression plots with age- and sex-adjusted axes display the best-fit line with 95% confidence intervals and each subject as a data point. Regression coefficients and FDR-corrected *q* values are given with each plot. Statistical significance is denoted as n.s. (not significant), \**q* ≤ 0.05, \*\**q* ≤ 0.01, \*\*\**q* ≤ 0.001.

The whole SN and matrix reference estimates indicated greater QSM asymmetry with greater motor asymmetry, but neither association was significant. N1 and N4 retained their direction and statistical significance in every leave-one-out jackknife analysis, while additional covariate adjustment changed the estimates little (Supplementary Table 14 and Supplementary Fig. 8).

R2* and R1 asymmetry estimates generally pointed in the same direction as the QSM estimates in N1, N2 and N4, but none survived FDR correction (R2*: *q* = 0.234 for N1, N2 and N4; R1: *q* = 0.840 for N1, N2 and N4) (Supplementary Table 14 and Supplementary Fig. 7).

## Discussion

In this study, we investigated the differential vulnerability of the nigrosomes in vivo, its clinical correlates, and the qMRI signatures associated with it, within a degeneration-independent anatomical framework built from histology-derived masks of nigrosomes N1–N5 and multimodal 7 T qMRI. Nigrosomes differed from the surrounding SN matrix in tissue composition and disease involvement, with the largest changes in N1, N2 and N4 and a rostrocaudal gradient within the matrix. These findings recover in vivo the spatial pattern established by histology^7,9^. QSM asymmetry in N1 and N4 also tracked contralateral MDS-UPDRS III asymmetry, extending the framework from group-level topography to clinically relevant lateralization.

### Differential nigral vulnerability in vivo

We find non-uniform involvement within the SN with the largest pathological changes centred around the nigrosomes, in agreement with histological studies.^4,5,7,9^ The nigrosomes showed a tissue composition distinct from that of the surrounding matrix. PD then altered the relationship between these compartments, largely through changes in N1, N2 and N4. Notably, these regional differences were detectable even though the whole-substantia-nigra analysis showed no significant group effect, demonstrating the value of a subregional framework. The direction of effect for the vulnerable nigrosomes (N1, N2 and N4) is preserved and the effect sizes are inflated after referencing to the surrounding matrix, with N1 and N4 significant in all three parameters and N2 in quantitative susceptibility under both schemes. However, as internal referencing reduces between-subject variance and inflates effect sizes by design, the larger effect sizes are not evidence of a larger biological effect; they reflect the reduced between-subject variance of the referenced measure.

The finding of nigrosomal change concentrated around N1, N2, and N4 corresponds well with previous imaging literature showing clusters of disease-related contrast located in areas corresponding to N1, N2, and N4 in both quantitative^15,39–42^ and qualitative^16,18,43–45^ MRI sequences.

N3 and N5 showed no significant group differences, but limited power precludes interpreting this as definitive sparing. Both regions are small, contain few voxels after mask registration, and are particularly susceptible to partial-volume effects. The opposite-direction differences in N3 (R2*) and N5 (QSM) may therefore reflect a partial-volume exchange with the surrounding matrix rather than a genuine biological reversal.

The matrix also showed spatially non-uniform involvement. QSM and R1 displayed a rostrocaudal gradient, with the largest PD-control differences towards the caudal end, mirroring the cell-loss framework established histologically.^9^ R2* showed a dorsoventral gradient, with the PD-control difference emerging at the level of the ventral tier^21^, and increasing dorsally, within the tiers to which histology localises dopamine-containing cell loss.^9^ The rostrocaudal pattern is consistent with previous QSM studies of the whole SN^46^ and resembles multimodal qMRI gradients along the putamen.^47^ Together, these findings align with the preferential degeneration of the posterior nigrostriatal system and its clinical expression in PD.

Note, however, that the matrix ROI is compositionally heterogeneous, as set out in the methods: its signal is a mixture in which only the contribution of the non-nigrosomal dopamine-containing neurons and glial cells are expected to change with disease. Matrix group differences and gradients are therefore conservative estimates, and the absence of a whole-matrix group effect should not be read as sparing of the dopamine-containing neurons within the matrix. Consistent with this, the R2* difference along the dorsoventral axis was close to zero in the most ventral bins, which correspond largely to the pars reticulata, and emerged only in the more dorsal bins containing the non-nigrosomal dopamine-containing neurons.

### Neurodegenerative processes altering qMRI contrast

Iron and myelin are the principal tissue determinants of the qMRI contrasts measured here. Iron raises QSM, R2* and R1. Tissue macromolecules, particularly myelin, raise R2* and R1 but lower QSM. This sign asymmetry is what allows the three contrasts, taken jointly, to be interpreted rather than merely detected, in terms of the pathological alterations responsible for a given change.^48^ We interpret our findings in light of four processes acting on this substrate, two primary and two secondary, while stressing that qMRI cannot separately quantify their contributions and that this reconstruction is inferential rather than demonstrated here. Early in disease, while dopamine-containing neurons are still present, iron accumulating in the pigment neuromelanin within them would be expected to raise all three quantitative parameters in the nigrosomes, as compared to surrounding matrix;^11^ as neurons die and microglia clear their neuromelanin-bound iron, this contribution will decrease. Secondary microglial activation and microglial invasion together with oligodendroglial and astrocytic scarring around degenerating neurons^49^ would raise local iron, as glial cells are iron rich^48^ and again increase QSM, R2* and R1, whereas cell death and loss of the efferent fibres leaving the nigrosomes would reduce myelin and other macromolecules, lowering R2* and R1 while raising QSM.

### qMRI contrast detects secondary processes rather than cell loss directly

Applied to our findings: in HCs, higher QSM and lower R1 in the nigrosomes than the matrix are consistent with the higher iron content of dopamine-containing neurons and low myelin content in SN.^11^ In patients with PD, this pattern widens - the nigrosome-matrix gap increases for QSM and R2* and narrows for R1 – predominantly through changes within the nigrosome rather than the matrix in every case. Because most nigral dopamine-containing neurons are already lost by the time of clinical motor onset, a cohort at this disease stage is better positioned to detect the secondary, iron-accumulating processes than the early, neuron-intrinsic ones, and we interpret the widened gap accordingly as predominantly secondary rather than a direct signature of ongoing neuronal iron handling.

The overall pattern - increases in patients across all three quantitative parameters within the nigrosomes despite the neuronal and myelin loss, and the gradients observed in all three modalities – suggests that secondary processes, in particular iron accumulation associated with inflammation and gliosis, may make a substantial contribution to the observed MRI contrast changes in PD. This interpretation is consistent with histological evidence showing that, in the healthy brain, iron contained in dopamine-containing neurons accounts for only 11–30% of the total iron within the nigrosomes.^11^ In PD, cellular iron concentration has been reported to increase by approximately 60% when averaged across all cell populations, including glial cells.^48^ Thus, iron accumulation associated with secondary cellular processes may outweigh the reduction in iron resulting from the loss of dopamine-containing neurons, potentially making a substantial contribution to the changes observed in qMRI parameters. This should be read as consistent with, rather than proof of, iron-dominated secondary pathology;^48^ per-nigrosome histological validation using advanced iron quantification and biophysical modelling would be needed to establish it directly.

### Relation to the histological ordering of cell loss

In the original publication by Damier and colleagues, a decrease in dopamine-containing neurons within nigrosomes was found with increasing disease duration, and over the five patients studied, N1 had the most degeneration, followed by N2, N4, N3, and N5.^9^ In our data, deviations from the healthy control mean across disease tertiles were most pronounced in N4, N1, and N2, with a tendency towards larger deviations in the longer duration tertiles. However, as none of these duration slopes survived correction, this observation is descriptive only. The ordering was not formally tested by pairwise comparison between nigrosomes, and effect sizes and z-scores reflect regional variability and partial-volume effects as well as biological change. Moreover, without trajectories for modelling neurodegeneration from qMRI signal, we cannot infer any temporal order of cell loss. We note too that the fine sequence reported post-mortem was itself inferred cross-sectionally from a small number (n=5) of late-stage cases without pairwise testing; the finding our data support is the gross vulnerability gradient, not a specific temporal order. Some MRI studies have reported correlations between signal and disease progression leading with N1 abnormality; however, these studies rely on the binary visual assessment of normal/abnormal nigrosomes and thus fail to capture the dynamics of neurodegeneration.^16,43,44^

### Clinical relevance of nigrosomal asymmetry

We find robust significant correlations between asymmetry in QSM for N1 and N4 and residual contralateral MDS-UPDRS III scores, with N1 showing the strongest correlation. Because the comparison is within-subject, asymmetry in secondary tissue alteration within the nigrosomes may track the lateralisation and severity of disease more faithfully than an absolute measure. The effects seem highly localized, with no other nigrosomes showing correlations (Supplementary Table 14).

At 7 T, correlations between asymmetry in R2* signal in the DNH and contralateral MDS-UPDRS III have previously been reported.^15^ We reproduce this association in QSM but not in R2*. A likely contributor is the discrepancy in the definition of the anatomical areas, as the DNH is defined as an area of low-susceptibility, low-R2* contrast, whereas ours is defined from calbindin-based histological masks independent of the qMRI contrast. Because N1 does not coincide anatomically with DNH,^14^ cross-cohort discrepancies of this kind are expected, and a shared, histology-anchored pipeline would be needed to resolve them. At 3 T, visual assessment of the DNH shows good agreement with contralateral clinical laterality,^50,51^ consistent with the lateralized susceptibility signal we observe.

### Limitations

The principal limitation of this study is that it rests on a single cohort with no independent replication and no per-subject histological confirmation; the entire inference depends on warping post-mortem masks into one cohort template, so registration accuracy and partial-volume effects are the dominant sources of uncertainty. In addition, the control group was drawn from two sources (nine STRAT-PARK and 19 quality-assurance participants); although mean SN signal did not differ between them (Supplementary Table 4), a comparison of nine versus 19 participants cannot establish equivalence, and residual cohort effects cannot be excluded.

Unlike in histological staining where the nigrosomes are clearly visible per subject, the nigrosomes are not reliably identifiable by in vivo MRI due to complex and partly unknown contrast mechanisms, small size and complex geometry. This is for example demonstrated by poor inter and intra-rater reliability in attempts to identify nigrosomes in MRI images.^18^ This study assumes that the nigrosomes are approximately similar in morphology and at approximately the same locations within the SN across individuals. It then relies on the template creation and focused atlas registration to capture the full nigrosomal morphology. Placing atlas-defined nigrosome masks in patients rests on two assumptions. The first, that the mutual arrangement of nigrosomes is preserved in PD, is supported by histology.^9^ The second is harder to verify: the warps that placed the masks in each participant were driven by the same quantitative contrasts in which the disease effect was measured, so disease-related changes in nigral contrast could in principle shift the masks systematically in patients relative to controls. We consider this unlikely, as the effect held at the most conservative mask thresholds, which confine each mask to its core, and the whole-nigra ROI, subject to identical warps, showed no group difference. Definitive validation would require post-mortem PD brains imaged with the same contrasts and stained for calbindin, which are not yet available. Beyond this, for smaller nigrosomes such as N3 and N5, partial volume effects, misregistration, and low power due to the small number of voxels and subjects make them more exposed to false negatives. For the larger nigrosomes, close proximities between nigrosomes may cause false positive results from partial volume effects and misregistration.

While being able to detect secondary degeneration processes with qMRI contrasts, QSM and R2* were demonstrated to be limited by not being able to directly detect neurodegeneration. The complex biological processes and complex dynamics of qMRI change in neurodegeneration limit the translation of qMRI signal into the degree of cell loss, unlike the histological gold standard of cell counting.^9^ Longitudinal studies investigating neurodegeneration so far yield inconclusive results on changes in qMRI signal with disease duration^52^, and normative trajectories suggest they may not necessarily be linear.^53^ The QSM and R2* values are therefore not able to directly quantify the degree of cell loss, limiting an exact in vivo degeneration framework. A cross-sectional design and lack of prodromal subjects fail to properly capture the stages and temporal aspects of qMRI changes in PD. Our study relies on disease duration as a temporal proxy to investigate the temporal aspects of nigrosomal degeneration. However, this measure is suboptimal due to degeneration happening at different rates and inherent difficulties in defining disease onset in PD.^54–56^ Additionally, at the time of motor symptom onset, most of the dopamine-containing cells are already lost, especially in the nigrosomes, hence producing little detectable signal change due to cell loss.

Lastly, the assessment of patients in the “ON” dopaminergic state reduces sensitivity and may dilute the MDS-UPDRS III asymmetry scores. Levodopa acts preferentially on the most affected side, hence compressing asymmetry in the “ON” state^57^. Striatal dopaminergic denervation indeed correlates most strongly with motor severity in the “OFF” state.^58^

Notwithstanding these limitations, a histology-anchored, degeneration-independent nigrosome framework recovered, in a modest and clinically heterogeneous cohort, a spatial pattern of vulnerability consistent with post-mortem findings across three complementary quantitative contrasts, two referencing schemes and a range of mask thresholds; we consider this robustness the principal strength of the approach.

### Future directions

Several extensions follow directly from the limitations of this work. The most immediate is validation in independent cohorts, and extension to conventional field strength, to establish that a histology-anchored nigrosome framework generalises beyond a single 7 T sample and is portable to the 3 T systems used in most clinical and research settings.

Extending the iron-sensitive contrasts used here with additional sequences would also give more information for resolving the complex dynamics of neurodegeneration. Recent biophysical modelling work suggests that another qMRI parameter, R2’, may provide higher specificity for neuronal loss, as it does not merely reflect total iron content, as QSM does, but is more sensitive to iron located in large cells, such as dopamine-containing neurons.^11^ This hypothesis, however, remains to be tested in clinical settings. Including a neuromelanin-sensitive sequence could also give an additional measure of degeneration of dopamine-containing cells, complementing the iron signal investigated in this study. Indeed, studies have shown a tight relationship between iron accumulation and loss of neuromelanin signal with PD progression.^15,59^ In parallel, advancements in biophysical modelling that separates the iron and myelin contributions to the measured relaxation and susceptibility^60^ could move the readout from where change occurs toward what it reflects, and, combined with per-region calibration, may eventually permit estimates of neuronal loss rather than of its secondary signature.

Moving from a spatial to a genuinely temporal account will require designs this study could not provide. Prodromal cohorts, including individuals with isolated REM-sleep behaviour disorder, could capture the earliest phase in which dopamine-containing neurons are still present and their loss potentially detectable, while longitudinal follow-up would allow subregional trajectories to be estimated within individuals rather than inferred cross-sectionally from disease duration.

The framework established here also invites more targeted questions. A spatial decomposition of the dorsolateral nigral hyperintensity would test whether its signal reflects N1 cell bodies together with their efferent projections, clarifying the long-standing conflation of this imaging sign with N1. Extending the approach to atypical parkinsonian syndromes would establish whether subregional signatures carry differential-diagnostic value. And because the nigrosome-matrix organisation is the nigral analogue of the striosome–matrix mosaic of the striatum, a shared, histology-anchored treatment of both compartments may offer a route to studying selective vulnerability across the nigrostriatal system as a whole.

## Data availability

The nigrosome and SN probability masks used in this study are publicly available (doi: 10.1038/s41597-026-08133-y).^21^ The cohort-specific multimodal templates and the region-of-interest values underlying all figures are provided as Source Data and will be deposited in Zenodo upon publication. Individual-level raw MRI data are not publicly available because their use is governed by participant consent, applicable data-protection legislation, and the regional ethics approvals for STRAT-PARK (REK 74985) and additional healthy controls (REK 108066). Access requests to the data pertaining to STRAT-PARK may be submitted to the STRAT-PARK consortium (K.E.S., C.T.) and will be reviewed by the STRAT-PARK leadership. Access requests to the additional healthy controls may be submitted to E.M.B. Access may be granted for scientifically and ethically approved purposes, subject to the necessary institutional approvals and data-sharing agreements.

## Code availability

Code used for MRI postprocessing and statistical analysis is publicly available at https://github.com/clinical-brain-systems/qMRI_Nigrosome_vulnerability. The script used for focused registration of the nigrosome atlas^21^ is available at https://osf.io/gsphy and was used without modification.

## Acknowledgements

We are deeply grateful to the study participants and their families involved in the study for their unique contribution. Furthermore, we would like to thank the Norwegian 7 T MR Center, especially medical physicist Maria Tunset Grinde, senior engineer Christopher Trimble, and senior radiographer Knut Kristian Nordlid. We are also grateful to nurse Anne Grete Wahlvåg for patient recruitment logistics, and MRI physicist Marc-Antoine Fortin for the help provided with establishing the post-processing qMRI pipeline used in the current study.

## Supplementary Information

See supplementary information file.

## Author contributions

**Ruiyi Li**: Conceptualization, Methodology, Investigation, Formal Analysis, Software, Visualization, and Writing (original draft, review & editing). **Evgeniya Kirilina**: Conceptualization, Methodology, Investigation, Formal Analysis, and Writing (review & editing). **Pierre-Louis Bazin**: Methodology, Investigation, Formal Analysis, Software, and Writing (review & editing). **Kjersti Eline Stige**: Investigation, Resources, and Writing (review & editing). **Runa G. Unsgård**: Resources, and Writing (review & editing). **Malte Brammerloh**: Software, Visualization, and Writing (review & editing). **Ingrid Gylterud Kvålsgard**: Resources, and Writing (review & editing). **Anne Louise Kristoffersen**: Resources, Methodology, and Writing (review & editing). **Erik M. Berntsen**: Resources and Writing (review & editing). **Pål Erik Goa**: Methodology, Supervision, Project administration, Funding acquisition, and Writing (review & editing). **Philippe Damier**: Conceptualization, Methodology, Investigation, and Writing (review & editing). **Charalampos Tzoulis**: Clinical Investigation, Resources, Supervision, Project Administration, Funding Acquisition, and Writing (Review and Editing). **Thanh P. Doan**: Conceptualization, Methodology, Investigation, Supervision, Funding Acquisition, and Writing (original draft, review & editing).

## References

1 Dorsey, E. R., Sherer, T., Okun, M. S. & Bloem, B. R. The Emerging Evidence of the Parkinson Pandemic. J Parkinsons Dis 8, S3–s8 (2018). 10.3233/jpd-181474

2 Welter, L. et al. The Costs of Parkinson’s Disease in Europe: Results of the Costs of Illness in Neurology Initiative. Mov Disord (2026). 10.1002/mds.70491

3 McGregor, M. M. & Nelson, A. B. Circuit Mechanisms of Parkinson’s Disease. Neuron 101, 1042–1056 (2019). 10.1016/j.neuron.2019.03.004

4 Tretiakoff, C. Contribution a l’etude de l’anatomie pathologique du locus niger de Soemmering avec quelques deductions relatives a la pathogenie des troubles du tonus musculaire et de la maladie de Parkinson. Thèse de Paris No. 293 (1919).

5 Hassler, R. Zur pathologischen Anatomie des parkinsonismus und der postenzephalitischen Parkinsonkrankheit. . Journal für Psychologie und Neurologie 48, 387–476 (1938).

6 Fearnley, J. M. & Lees, A. J. Ageing and Parkinson’s disease: substantia nigra regional selectivity. Brain 114 ( Pt 5), 2283–2301 (1991). 10.1093/brain/114.5.2283

7 Damier, P., Hirsch, E. C., Agid, Y. & Graybiel, A. M. The substantia nigra of the human brain. I. Nigrosomes and the nigral matrix, a compartmental organization based on calbindin D(28K) immunohistochemistry. Brain 122 (Pt 8), 1421–1436 (1999). 10.1093/brain/122.8.1421

8 Graybiel, A. M. & Ragsdale, C. W., Jr. Histochemically distinct compartments in the striatum of human, monkeys, and cat demonstrated by acetylthiocholinesterase staining. Proc Natl Acad Sci U S A 75, 5723–5726 (1978). 10.1073/pnas.75.11.5723

9 Damier, P., Hirsch, E. C., Agid, Y. & Graybiel, A. M. The substantia nigra of the human brain. II. Patterns of loss of dopamine-containing neurons in Parkinson’s disease. Brain 122 (Pt 8), 1437–1448 (1999). 10.1093/brain/122.8.1437

10 Halliday, G., Reyes, S. & Double, K. in The Human Nervous System (Third Edition) (eds Jürgen K. Mai & George Paxinos) 439–455 (Academic Press, 2012).

11 Brammerloh, M. et al. Measuring the iron content of dopaminergic neurons in substantia nigra with MRI relaxometry. Neuroimage 239, 118255 (2021). 10.1016/j.neuroimage.2021.118255

12 Blazejewska, A. I. et al. Visualization of nigrosome 1 and its loss in PD: pathoanatomical correlation and in vivo 7 T MRI. Neurology 81, 534–540 (2013). 10.1212/WNL.0b013e31829e6fd2

13 Lee, H., Baek, S. Y., Chun, S. Y., Lee, J. H. & Cho, H. Specific visualization of neuromelanin-iron complex and ferric iron in the human post-mortem substantia nigra using MR relaxometry at 7T. Neuroimage 172, 874–885 (2018). 10.1016/j.neuroimage.2017.11.035

14 Brammerloh, M. et al. Swallow Tail Sign: Revisited. Radiology 305, 674–677 (2022). 10.1148/radiol.212696

15 Madelung, C. F. et al. High-resolution mapping of substantia nigra in Parkinson’s disease using 7 tesla magnetic resonance imaging. NPJ Parkinsons Dis 11, 113 (2025). 10.1038/s41531-025-00972-7

16 Shin, N. Y. et al. Cortical thinning pattern according to differential nigrosome involvement in patients with Parkinson’s disease. NeuroImage: Clinical 28 (2020). 10.1016/j.nicl.2020.102382

17 Kim, E. Y., Sung, Y. H. & Lee, J. Nigrosome 1 imaging: technical considerations and clinical applications. Br J Radiol 92, 20180842 (2019). 10.1259/bjr.20180842

18 Schwarz, S. T. et al. Parkinson’s disease related signal change in the nigrosomes 1–5 and the substantia nigra using T2* weighted 7T MRI. NeuroImage: Clinical 19, 683–689 (2018). 10.1016/j.nicl.2018.05.027

19 Weiskopf, N., Edwards, L. J., Helms, G., Mohammadi, S. & Kirilina, E. Quantitative magnetic resonance imaging of brain anatomy and in vivo histology. Nature Reviews Physics 3, 570–588 (2021). 10.1038/s42254-021-00326-1

20 Edwards, L. J. et al. Quantitative MRI maps of human neocortex explored using cell type-specific gene expression analysis. Cerebral Cortex 33, 5704–5716 (2023). 10.1093/cercor/bhac453

21 Brammerloh, M. et al. A neuroimaging atlas of the nigrosomes in the substantia nigra based on 3D histology. Sci Data 13 (2026). 10.1038/s41597-026-08133-y

22 Stige, K. E. et al. The STRAT-PARK cohort: A personalized initiative to stratify Parkinson’s disease. Prog Neurobiol 236, 102603 (2024). 10.1016/j.pneurobio.2024.102603

23 Postuma, R. B. et al. MDS clinical diagnostic criteria for Parkinson’s disease. Mov Disord 30, 1591–1601 (2015). 10.1002/mds.26424

24 Unsgård, R. G. et al. 7 Tesla MRI with Quantitative Susceptibility Mapping: qualitative grading and quantitative focal measurements of the substantia nigra as a potential biomarker for Parkinson’s disease. Neuroradiology (2026). 10.1007/s00234-026-04133-2

25 Marques, J. P. et al. MP2RAGE, a self bias-field corrected sequence for improved segmentation and T1-mapping at high field. Neuroimage 49, 1271–1281 (2010). 10.1016/j.neuroimage.2009.10.002

26 Eckstein, K. et al. Computationally Efficient Combination of Multi-channel Phase Data From Multi-echo Acquisitions (ASPIRE). Magn Reson Med 79, 2996–3006 (2018). 10.1002/mrm.26963

27 Chung, S., Kim, D., Breton, E. & Axel, L. Rapid B1+ mapping using a preconditioning RF pulse with TurboFLASH readout. Magn Reson Med 64, 439– 446 (2010). 10.1002/mrm.22423

28 Huntenburg, J. M., Steele, C. J. & Bazin, P. L. Nighres: processing tools for high-resolution neuroimaging. Gigascience 7 (2018). 10.1093/gigascience/giy082

29 Bazin, P. L. et al. Denoising High-Field Multi-Dimensional MRI With Local Complex PCA. Front Neurosci 13, 1066 (2019). 10.3389/fnins.2019.01066

30 Tustison, N. J. et al. N4ITK: improved N3 bias correction. IEEE Trans Med Imaging 29, 1310–1320 (2010). 10.1109/tmi.2010.2046908

31 Langkammer, C. et al. Fast quantitative susceptibility mapping using 3D EPI and total generalized variation. Neuroimage 111, 622–630 (2015). 10.1016/j.neuroimage.2015.02.041

32 Tustison, N. J. et al. The ANTsX ecosystem for quantitative biological and medical imaging. Sci Rep 11, 9068 (2021). 10.1038/s41598-021-87564-6

33 Lancione, M. et al. High resolution multi-parametric probabilistic in vivo atlas of dorsolateral nigral hyperintensity via 7 T MRI. Sci Data 12, 958 (2025). 10.1038/s41597-025-05325-w

34 Amunts, K. et al. BigBrain: an ultrahigh-resolution 3D human brain model. Science 340, 1472–1475 (2013). 10.1126/science.1235381

35 Makris, N. et al. Decreased volume of left and total anterior insular lobule in schizophrenia. Schizophr Res 83, 155–171 (2006). 10.1016/j.schres.2005.11.020

36 Desikan, R. S. et al. An automated labeling system for subdividing the human cerebral cortex on MRI scans into gyral based regions of interest. Neuroimage 31, 968–980 (2006). 10.1016/j.neuroimage.2006.01.021

37 Frazier, J. A. et al. Structural brain magnetic resonance imaging of limbic and thalamic volumes in pediatric bipolar disorder. Am J Psychiatry 162, 1256–1265 (2005). 10.1176/appi.ajp.162.7.1256

38 Goldstein, J. M. et al. Hypothalamic abnormalities in schizophrenia: sex effects and genetic vulnerability. Biol Psychiatry 61, 935–945 (2007). 10.1016/j.biopsych.2006.06.027

39 Seong, M., Park, S., Sung, Y. H. & Kim, E. Y. Diagnostic performance of a high-spatial-resolution voxelwise analysis of neuromelanin-sensitive imaging in early-stage idiopathic Parkinson’s disease. BMC Med Imaging 23, 64 (2023). 10.1186/s12880-023-01018-1

40 Park, S., Sung, Y. H., Kim, W. R., Noh, Y. & Kim, E. Y. Correlation Between Neuromelanin-Sensitive MRI and18F-FP-CIT PET in Early-Stage Parkinson’s Disease: Utility of a Voxel-Wise Analysis by Using High-Spatial-Resolution MRI. Journal of Clinical Neurology (Korea) 19, 156–164 (2023). 10.3988/jcn.2022.0147

41 Sung, Y. H., Noh, Y. & Kim, E. Y. Early-stage Parkinson’s disease: Abnormal nigrosome 1 and 2 revealed by a voxelwise analysis of neuromelanin-sensitive MRI. Human Brain Mapping 42, 2823–2832 (2021). 10.1002/hbm.25406

42 Langley, J. et al. Multimodal assessment of nigrosomal degeneration in Parkinson’s disease. Parkinsonism Relat Disord 80, 102–107 (2020). 10.1016/j.parkreldis.2020.09.021

43 Yeop Kim, E., Noh, Y., Sung, Y. H., Lee, J. & Seo, S. In vivo nigrosome 1 and 4 imaging at 3T and Its Correlation with Dopamine Transporter imaging in Idiopathic Parkinson’s Disease. Neuroradiology 60, 108–109 (2018).

44 Sung, Y. H. et al. Differential involvement of nigral subregions in idiopathic parkinson’s disease. Human Brain Mapping 39, 542–553 (2018). 10.1002/hbm.23863

45 Kim, E. Y. & Sung, Y. H. Early versus late stage idiopathic parkinson’s disease differential involvement of nigrosome 1 and 4. Neuroradiology 59, S72–S73 (2017). 10.1007/s00234-017-1872-5

46 Lotfipour, A. K. et al. High resolution magnetic susceptibility mapping of the substantia nigra in Parkinson’s disease. J Magn Reson Imaging 35, 48–55 (2012). 10.1002/jmri.22752

47 Drori, E., Cohen, L., Arkadir, D., Yahalom, G. & Mezer, A. A. Multiparametric quantitative MRI uncovers putamen microstructural changes in Parkinson’s disease. NPJ Parkinsons Dis 11, 197 (2025). 10.1038/s41531-025-01020-0

48 Friedrich, I. et al. Cell specific quantitative iron mapping on brain slices by immuno-µPIXE in healthy elderly and Parkinson’s disease. Acta Neuropathol Commun 9, 47 (2021). 10.1186/s40478-021-01145-2

49 Damier, P., Hirsch, E. C., Zhang, P., Agid, Y. & Javoy-Agid, F. Glutathione peroxidase, glial cells and Parkinson’s disease. Neuroscience 52, 1–6 (1993). 10.1016/0306-4522(93)90175-f

50 Stezin, A. et al. Clinical utility of visualisation of nigrosome-1 in patients with Parkinson’s disease. Eur Radiol 28, 718–726 (2018). 10.1007/s00330-017-4950-5

51 Noh, Y., Sung, Y. H., Lee, J. & Kim, E. Y. Nigrosome 1 Detection at 3T MRI for the Diagnosis of Early-Stage Idiopathic Parkinson Disease: Assessment of Diagnostic Accuracy and Agreement on Imaging Asymmetry and Clinical Laterality. AJNR Am J Neuroradiol 36, 2010–2016 (2015). 10.3174/ajnr.A4412

52 Guan, X. et al. Neuroimaging of Parkinson’s disease by quantitative susceptibility mapping. Neuroimage 289, 120547 (2024). 10.1016/j.neuroimage.2024.120547

53 Chen, X. et al. Mapping heterogeneous region- and tissue-specific brain ageing patterns using quantitative MRI. Brain Commun 8, fcag010 (2026). 10.1093/braincomms/fcag010

54 Dadu, A. et al. Identification and prediction of Parkinson’s disease subtypes and progression using machine learning in two cohorts. NPJ Parkinsons Dis 8, 172 (2022). 10.1038/s41531-022-00439-z

55 Hähnel, T. et al. Progression subtypes in Parkinson’s disease identified by a data-driven multi cohort analysis. NPJ Parkinsons Dis 10, 95 (2024). 10.1038/s41531-024-00712-3

56 Fraza, C. et al. Understanding individual neurodegenerative progression in Parkinson’s disease through normative modelling. Sci Rep 15, 36659 (2025). 10.1038/s41598-025-20484-x

57 Martinu, K., Nagano-Saito, A., Fogel, S. & Monchi, O. Asymmetrical effect of levodopa on the neural activity of motor regions in PD. PLoS One 9, e111600 (2014). 10.1371/journal.pone.0111600

58 Pirker, W. Correlation of dopamine transporter imaging with parkinsonian motor handicap: how close is it? Mov Disord 18 Suppl 7, S43–51 (2003). 10.1002/mds.10579

59 Biondetti, E. et al. The spatiotemporal changes in dopamine, neuromelanin and iron characterizing Parkinson’s disease. Brain 144, 3114–3125 (2021). 10.1093/brain/awab191

60 Stüber, C. et al. Myelin and iron concentration in the human brain: a quantitative study of MRI contrast. Neuroimage 93 Pt 1, 95–106 (2014). 10.1016/j.neuroimage.2014.02.026

