## Supplemental Files for "Selective nigrosome vulnerability in Parkinson’s disease revealed by histology-anchored ultra-high-field quantitative MRI"

Contents

Supplementary Fig. 1 Multimodal quantitative MRI cohort-specific templates at the level of the substantia nigra
Supplementary Table 1 Probabilistic nigrosome mask diagnostics
Supplementary Fig. 2 Nigrosome mask probabilistic Dice overlap
Supplementary Table 2 Aggregated demographic statistics, patient cohort
Supplementary Table 3 Aggregated demographic statistics, control cohort
Supplementary Table 4 Between-control-cohort comparison of mean substantia nigra signal
Supplementary Table 5 Comparison of mean substantia nigra signal with versus without small vessel disease
Supplementary Table 6 Pooled nigrosome versus matrix, within-group gaps
Supplementary Table 7 Pooled nigrosome versus matrix gap, PD-versus-HC contrast
Supplementary Table 8 Pooled nigrosome versus matrix, region-separate PD-versus-HC
Supplementary Table 9 Per-nigrosome group differences, standard reference
Supplementary Table 10 Per-nigrosome group differences, matrix reference
Supplementary Fig. 3 Sweep of per-nigrosome effect across mask thresholds and modalities
Supplementary Table 11 Substantia nigra matrix gradient linear mixed models
Supplementary Fig. 4 Graphical summary of gradient interactions
Supplementary Table 12 Nigrosome signal and disease duration, standard reference
Supplementary Table 13 Nigrosome signal and disease duration, matrix reference
Supplementary Fig. 5 Deviation from the healthy control distribution by disease-duration tertile
Supplementary Fig. 6 Deviation from the healthy control distribution per participant
Supplementary Table 14 Imaging asymmetry and contralateral clinical motor asymmetry
Supplementary Fig. 7 Motor asymmetry summary plots
Supplementary Fig. 8 Motor asymmetry, covariate sensitivity

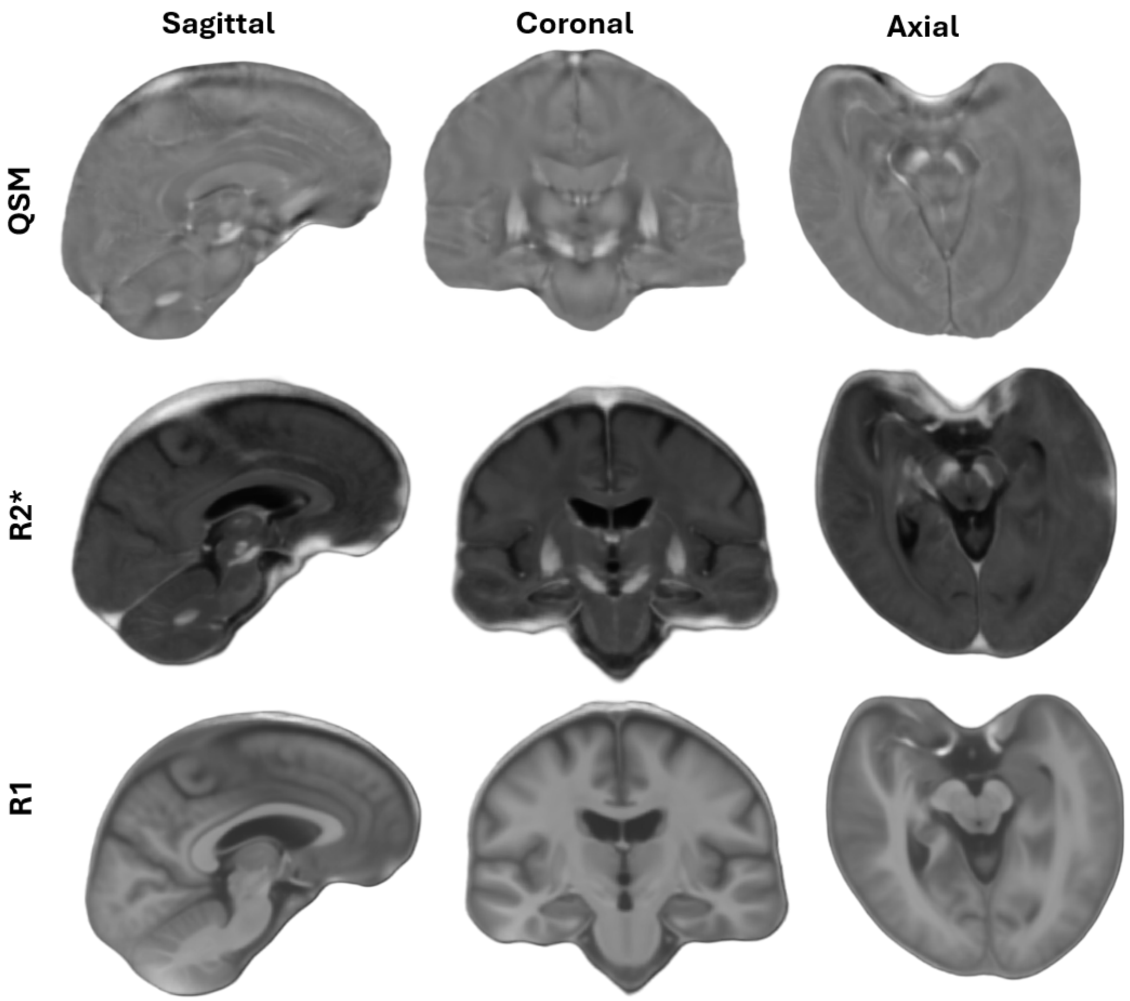


**Supplementary Fig. 1 |** Multimodal quantitative MRI (qMRI) cohort-specific templates at the level of the substantia nigra (SN). Sagittal, coronal, and axial views of the study-specific quantitative susceptibility mapping (QSM), R2*, and R1 templates constructed from all 56 participants (28 Parkinson's disease (PD), 28 healthy controls (HC)) with advanced normalization tools (ANTs) multivariate template construction. Slices are centred on the SN. Left–right convention is radiological.

**Supplementary Table 1 |** Probabilistic nigrosome mask diagnostics (bilateral masks; voxel volume 0.422 mm³).

| ROI | N voxels (p > 0) | Effective N (Kish) | Effective volume (mm³) |
|---|---|---|---|
| N1 | 1009 | 463.7 | 195.6 |
| N2 | 458 | 197.4 | 83.3 |
| N3 | 74 | 30.1 | 12.7 |
| N4 | 285 | 126.1 | 53.2 |
| N5 | 126 | 53.5 | 22.6 |
| SN | 13730 | 4129.2 | 1742.0 |
| matrix | 13712 | 4087.1 | 1724.2 |

Effective *n* is the Kish effective sample size, $(\Sigma w)^2/\Sigma w^2$, of the voxel probability weights $w$, and effective volume is effective *n* multiplied by voxel volume. N1–N5, nigrosomes 1–5; matrix, substantia nigra outside the nigrosomes.

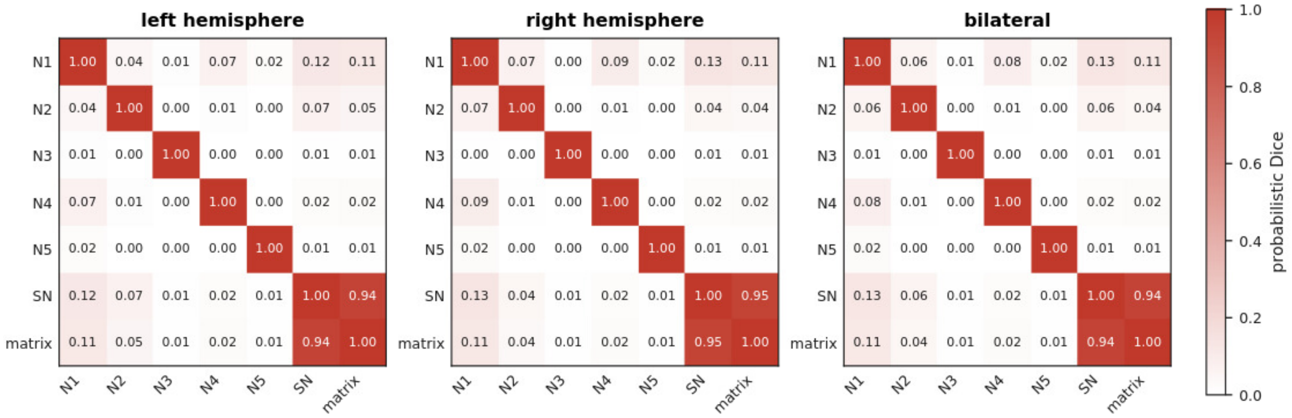


**Supplementary Fig. 2 |** Left, right, and bilateral probabilistic Dice overlap. Pairwise probabilistic dice overlap between the left, right, and bilateral nigrosome masks (N1–N5), the whole SN and the matrix. The near-zero overlap among N1–N5 confirms that the nigrosome masks are spatially disjoint.

**Supplementary Table 2 |** Aggregated demographic statistics for patient cohort (n = 28).

| Characteristic | Value |
|---|---|
| Age, years, mean (SD) | 66.8 (6.8) |
| Age, years, range | 51.0 to 80.0 |
| Sex, female, n (%) | 13 (46.4%) |
| Sex, male, n (%) | 15 (53.6%) |
| Disease duration, years, mean (SD) | 8.1 (5.8) |
| Disease duration, years, range | 1.3 to 28.2 |
| Hoehn and Yahr stage, mean (SD) | 2.0 (0.6) |
| Hoehn and Yahr stage, range | 1.0 to 3.0 |
| Hoehn and Yahr stage 1, n (%) | 6 (21.4%) |
| Hoehn and Yahr stage 2, n (%) | 17 (60.7%) |
| Hoehn and Yahr stage 3, n (%) | 5 (17.9%) |
| MDS-UPDRS part III total, mean (SD) | 25.3 (12.4) |
| MDS-UPDRS part III total, range | 9.0 to 51.0 |
| Clinically more affected side, right, n (%) [a] | 11 (39.3%) |
| Clinically more affected side, left, n (%) [a] | 14 (50.0%) |
| Clinically more affected side, symmetric, n (%) [a] | 3 (10.7%) |
| LEDD, mg/day, mean (SD) | 652 (386) |
| LEDD, mg/day, range | 300 to 1780 |
| Time since last levodopa dose, min, mean (SD) | 127 (58) |
| Time since last levodopa dose, min, range | 15 to 225 |
| Levodopa treatment, n (%) | 27 (96.4%) |

[a]Defined from the lateralised MDS-UPDRS part III items; symmetric when the right and left subscores were equal. LEDD, levodopa-equivalent daily dose; MDS-UPDRS, Movement Disorder Society Unified Parkinson’s Disease Rating Scale; SD, standard deviation. MDS-UPDRS part III and Hoehn and Yahr stage were assessed in the “ON” dopaminergic state.

**Supplementary Table 3 |** Aggregated demographic statistics for control cohort (n = 28).

| Characteristic | Value |
|---|---|
| Age, years, mean (SD) | 66.0 (8.6) |
| Age, years, range | 45.0 to 80.0 |
| Sex, female, n (%) | 13 (46.4%) |
| Sex, male, n (%) | 15 (53.6%) |

Controls comprised nine participants recruited within STRAT-PARK and 19 from the quality-assurance cohort; the two sources are compared in Supplementary Table 4.

**Supplementary Table 4 |** Sensitivity analysis: between-control-cohort comparison of age- and sex-adjusted mean substantia nigra signal.

| Modality | *n* (STRAT-PARK / QA cohort) | Adjusted β [95% CI] | Permutation *P* | *q* (BH) |
|---|---|---|---|---|
| QSM | 9/19 | −0.0053 [−0.0216, 0.0077] | 0.500 | 0.750 |
| R2* | 9/19 | −1.28 [−10.79, 7.08] | 0.772 | 0.772 |
| R1 | 9/19 | −0.0139 [−0.0310, 0.0013] | 0.111 | 0.334 |

Healthy controls recruited within STRAT-PARK (n = 9) were compared with controls from the quality-assurance (QA) cohort (*n* = 19) on mean whole-SN signal, adjusting for age and sex. β is the STRAT-PARK-minus-QA difference in ppm (QSM) or $s^{-1}$ (R2*, R1) with bootstrap 95% confidence interval (CI). *P* from Freedman–Lane permutation; *q* from Benjamini–Hochberg (BH) correction across the three modalities.

**Supplementary Table 5 |** Sensitivity analysis: within-cohort comparison of age- and sex-adjusted mean substantia nigra signal in participants with versus without small vessel disease.

| Modality | *n* (small vessel disease / remainder) | Adjusted β [95% CI] | Permutation *P* | *q* (BH) |
|---|---|---|---|---|
| QSM | 4/15 | −0.0160 [−0.0408, 0.0143] | 0.227 | 0.423 |
| R2* | 4/15 | −8.68 [−21.43, 3.61] | 0.282 | 0.423 |
| R1 | 4/15 | 0.0011 [−0.0323, 0.0269] | 0.937 | 0.937 |

β is the difference (small vessel disease minus remainder) in mean whole-SN signal in ppm (QSM) or $s^{-1}$ (R2*, R1), adjusted for age and sex, with bootstrap 95% CI; *P* from Freedman–Lane permutation; *q* from BH correction across the three modalities. The controls with incidental small vessel disease were all older than 75 years with Fazekas grade 3.

**Supplementary Table 6 |** Pooled nigrosome vs matrix: within-group gaps.

| Modality | Group | Mean gap | SD gap | t | *P* | q (BH) |
|---|---|---|---|---|---|---|
| QSM | HC | 0.0234 | 0.0049 | 25.29 | $2.5 \times 10^{-20}$ | $7.4 \times 10^{-20}$ |
| QSM | PD | 0.0336 | 0.0079 | 22.42 | $5.5 \times 10^{-19}$ | $1.1 \times 10^{-18}$ |
| R2* | HC | 0.5960 | 2.8990 | 1.09 | 0.286 | 0.286 |
| R2* | PD | 3.3156 | 3.5039 | 5.01 | $3.0 \times 10^{-5}$ | $3.6 \times 10^{-5}$ |
| R1 | HC | -0.0318 | 0.0062 | -27.18 | $3.8 \times 10^{-21}$ | $2.3 \times 10^{-20}$ |
| R1 | PD | -0.0237 | 0.0058 | -21.85 | $1.1 \times 10^{-18}$ | $1.6 \times 10^{-18}$ |

Within-subject difference between the pooled nigrosome signal and the matrix signal (bilateral average), $n = 28$ per group. Units: ppm (QSM), $s^{-1}$ (R2*, R1). One-sample $t$-test against zero; $q$ from BH correction across the six tests.

**Supplementary Table 7 |** Pooled nigrosome vs matrix gap: PD-vs-HC contrast.

| Modality | Δ(PD-HC) | Bootstrap 95% CI | Hedges' g | Hedges' g 95% CI | Adjusted g | Adjusted g 95% CI | Perm *P* | q (BH) |
|---|---|---|---|---|---|---|---|---|
| QSM | 0.0101 | [0.0068, 0.0137] | 1.53 | [1.06, 2.15] | 1.52 | [1.05, 2.15] | <0.0001 | <0.0001 |
| R2* | 2.70 | [1.02, 4.35] | 0.83 | [0.32, 1.42] | 0.83 | [0.34, 1.44] | 0.003 | 0.003 |
| R1 | 0.0079 | [0.0051, 0.0110] | 1.33 | [0.92, 1.85] | 1.3 | [0.91, 1.78] | <0.0001 | <0.0001 |

Group difference in the nigrosome-minus-matrix gap ($n = 28$ PD, 28 HC), adjusted for age and sex. Units: ppm (QSM), $s^{-1}$ (R2*, R1). Adjusted Hedges' $g$ uses the covariate-adjusted group difference as numerator and the unadjusted pooled SD as denominator. $P$ from Freedman–Lane permutation; $q$ from BH correction across the three modalities.

**Supplementary Table 8 |** Pooled nigrosome vs matrix: region-separate PD-vs-HC.

| Modality | Region | β (PD-HC) | Bootstrap 95% CI | Hedges' g | Hedges' g 95% CI | Adjusted g | Adjusted g 95% CI | Perm *P* | q (BH) |
|---|---|---|---|---|---|---|---|---|---|
| QSM | Nigrosomes (pooled) | 0.0108 | [0.0025, 0.0188] | 0.69 | [0.19, 1.24] | 0.69 | [0.16, 1.27] | 0.011 | 0.017 |
| QSM | matrix | 0.0006 | [−0.0066, 0.0075] | 0.05 | [-0.5, 0.58] | 0.05 | [-0.5, 0.55] | 0.857 | 0.857 |
| R2* | Nigrosomes (pooled) | 5.22 | [0.61, 9.61] | 0.6 | [0.09, 1.2] | 0.6 | [0.06, 1.21] | 0.028 | 0.028 |
| R2* | matrix | 2.51 | [−1.18, 5.97] | 0.35 | [-0.16, 0.93] | 0.36 | [-0.16, 0.93] | 0.185 | 0.278 |
| R1 | Nigrosomes (pooled) | 0.0157 | [0.0059, 0.0251] | 0.73 | [0.22, 1.35] | 0.72 | [0.26, 1.29] | 0.003 | 0.008 |
| R1 | matrix | 0.0078 | [−0.0013, 0.0168] | 0.37 | [-0.14, 0.94] | 0.37 | [-0.06, 0.86] | 0.106 | 0.278 |

Age- and sex-adjusted group difference in mean signal ($n = 28$ PD, 28 HC). Units: ppm (QSM), $s^{-1}$ (R2*, R1). Adjusted Hedges' $g$ as in Supplementary Table 7. $P$ from Freedman–Lane permutation; $q$ from BH correction across the three modalities within each region.

**Supplementary Table 9 |** Per-nigrosome group differences (standard reference).

| Modality | ROI | β(PD-HC) | Bootstrap 95% CI | Hedges' g | Hedges' g 95% CI | Adjusted g | Adjusted g 95% CI | Perm *P* | q (BH) |
|---|---|---|---|---|---|---|---|---|---|
| QSM | N1 | 0.0101 | [0.0029, 0.0174] | 0.69 | [0.19, 1.25] | 0.69 | [0.17, 1.23] | 0.011 | 0.028 |
| QSM | N2 | 0.0136 | [0.0021, 0.0249] | 0.62 | [0.12, 1.22] | 0.61 | [0.1, 1.22] | 0.025 | 0.042 |
| QSM | N3 | −0.0034 | [−0.0120, 0.0044] | -0.21 | [-0.72, 0.32] | -0.21 | [-0.72, 0.31] | 0.424 | 0.424 |
| QSM | N4 | 0.0186 | [0.0097, 0.0272] | 1.09 | [0.6, 1.71] | 1.09 | [0.58, 1.74] | 0.0002 | 0.001 |
| QSM | N5 | −0.0041 | [−0.0118, 0.0040] | -0.27 | [-0.84, 0.24] | -0.26 | [-0.84, 0.24] | 0.326 | 0.408 |
| QSM | SN | 0.0017 | [−0.0055, 0.0088] | 0.12 | [-0.39, 0.66] | 0.12 | [-0.4, 0.65] | 0.647 | — |
| R2* | N1 | 5.66 | [1.11, 10.14] | 0.66 | [0.16, 1.28] | 0.66 | [0.1, 1.27] | 0.018 | 0.045 |
| R2* | N2 | 3.77 | [−2.06, 9.40] | 0.33 | [-0.18, 0.9] | 0.34 | [-0.21, 0.88] | 0.207 | 0.258 |
| R2* | N3 | −1.58 | [−6.19, 2.81] | -0.19 | [-0.73, 0.34] | -0.18 | [-0.72, 0.33] | 0.493 | 0.493 |
| R2* | N4 | 11.06 | [5.67, 16.47] | 1.07 | [0.56, 1.67] | 1.07 | [0.56, 1.7] | 0.0004 | 0.002 |
| R2* | N5 | 3.26 | [−0.74, 7.25] | 0.41 | [-0.12, 1.01] | 0.42 | [-0.09, 1.03] | 0.121 | 0.202 |
| R2* | SN | 2.80 | [−0.97, 6.34] | 0.39 | [-0.13, 0.99] | 0.39 | [-0.12, 1.0] | 0.147 | — |
| R1 | N1 | 0.0179 | [0.0077, 0.0275] | 0.82 | [0.31, 1.47] | 0.82 | [0.35, 1.39] | 0.001 | 0.003 |
| R1 | N2 | 0.0117 | [0.0005, 0.0225] | 0.49 | [-0.01, 1.09] | 0.48 | [0.01, 1.02] | 0.048 | 0.081 |
| R1 | N3 | 0.0016 | [−0.0097, 0.0126] | 0.05 | [-0.48, 0.59] | 0.06 | [-0.42, 0.52] | 0.784 | 0.784 |
| R1 | N4 | 0.0197 | [0.0097, 0.0297] | 0.9 | [0.38, 1.57] | 0.89 | [0.42, 1.51] | 0.0007 | 0.003 |
| R1 | N5 | 0.0034 | [−0.0054, 0.0124] | 0.16 | [-0.37, 0.71] | 0.17 | [-0.28, 0.64] | 0.466 | 0.582 |
| R1 | SN | 0.0086 | [−0.0004, 0.0178] | 0.41 | [-0.11, 0.97] | 0.41 | [-0.02, 0.91] | 0.075 | — |

Age- and sex-adjusted group difference in bilateral mean signal per region ($n = 28$ PD, 28 HC). Units: ppm (QSM), $s^{-1}$ (R2*, R1). Adjusted Hedges' *g* as in Supplementary Table 7. *P* from Freedman–Lane permutation; *q* from BH correction across N1–N5 within each modality; the whole-SN row is shown for context and was not included in the correction family. Dashes in the q column denote rows excluded from the correction family.

**Supplementary Table 10 |** Per-nigrosome group differences (matrix reference).

| Modality | ROI | β(PD-HC) | Bootstrap 95% CI | Hedges' g | Hedges' g 95% CI | Adjusted g | Adjusted g 95% CI | Perm *P* | q (BH) |
|---|---|---|---|---|---|---|---|---|---|
| QSM | N1 | 0.0094 | [0.0063, 0.0126] | 1.55 | [1.08, 2.14] | 1.56 | [1.08, 2.18] | <0.0001 | 0.0003 |
| QSM | N2 | 0.0129 | [0.0058, 0.0202] | 0.93 | [0.44, 1.54] | 0.9 | [0.43, 1.53] | 0.0008 | 0.001 |
| QSM | N3 | −0.0040 | [−0.0119, 0.0039] | -0.26 | [-0.77, 0.25] | -0.27 | [-0.81, 0.28] | 0.334 | 0.334 |
| QSM | N4 | 0.0179 | [0.0126, 0.0234] | 1.72 | [1.28, 2.33] | 1.72 | [1.29, 2.32] | <0.0001 | 0.0003 |
| QSM | N5 | −0.0047 | [−0.0092, −0.0005] | -0.56 | [-1.03, -0.09] | -0.55 | [-1.04, -0.07] | 0.040 | 0.050 |
| QSM | SN | 0.0011 | [0.0007, 0.0014] | 1.52 | [1.06, 2.18] | 1.51 | [1.05, 2.13] | <0.0001 | — |
| R2* | N1 | 3.15 | [1.06, 5.14] | 0.78 | [0.27, 1.41] | 0.77 | [0.27, 1.39] | 0.004 | 0.009 |
| R2* | N2 | 1.25 | [−1.77, 4.27] | 0.2 | [-0.32, 0.72] | 0.21 | [-0.34, 0.73] | 0.444 | 0.534 |
| R2* | N3 | −4.09 | [−7.46, −0.66] | -0.61 | [-1.17, -0.11] | -0.61 | [-1.16, -0.12] | 0.022 | 0.037 |
| R2* | N4 | 8.55 | [5.50, 11.71] | 1.38 | [0.85, 2.12] | 1.37 | [0.83, 2.12] | <0.0001 | 0.0005 |
| R2* | N5 | 0.75 | [−1.58, 3.06] | 0.16 | [-0.33, 0.76] | 0.17 | [-0.33, 0.78] | 0.534 | 0.534 |
| R2* | SN | 0.28 | [0.11, 0.45] | 0.83 | [0.33, 1.44] | 0.83 | [0.32, 1.42] | 0.003 | — |
| R1 | N1 | 0.0101 | [0.0063, 0.0139] | 1.37 | [0.96, 1.9] | 1.35 | [0.9, 1.91] | <0.0001 | 0.0003 |
| R1 | N2 | 0.0039 | [−0.0010, 0.0089] | 0.4 | [-0.12, 0.9] | 0.37 | [-0.1, 0.85] | 0.154 | 0.154 |
| R1 | N3 | −0.0062 | [−0.0143, 0.0007] | -0.44 | [-0.91, 0.06] | -0.42 | [-0.88, 0.08] | 0.118 | 0.154 |
| R1 | N4 | 0.0119 | [0.0079, 0.0163] | 1.32 | [0.88, 1.9] | 1.28 | [0.9, 1.82] | <0.0001 | 0.0003 |
| R1 | N5 | −0.0044 | [−0.0098, 0.0013] | -0.42 | [-0.95, 0.09] | -0.41 | [-0.93, 0.12] | 0.128 | 0.154 |
| R1 | SN | 0.0008 | [0.0005, 0.0012] | 1.35 | [0.92, 1.86] | 1.32 | [0.93, 1.81] | <0.0001 | — |

Each regional value was expressed relative to the subject's own matrix signal before group comparison by subtracting the mean matrix signal. Age- and sex-adjusted group difference in bilateral mean signal per region ($n$ = 28 PD, 28 HC). Units: ppm (QSM), $s^{-1}$ (R2*, R1). Adjusted Hedges' $g$ as in Supplementary Table 7. $P$ from Freedman–Lane permutation; $q$ from BH correction across N1–N5 within each modality; the whole-SN row is shown for context and was not included in the correction family. Dashes in the q column denote rows excluded from the correction family.

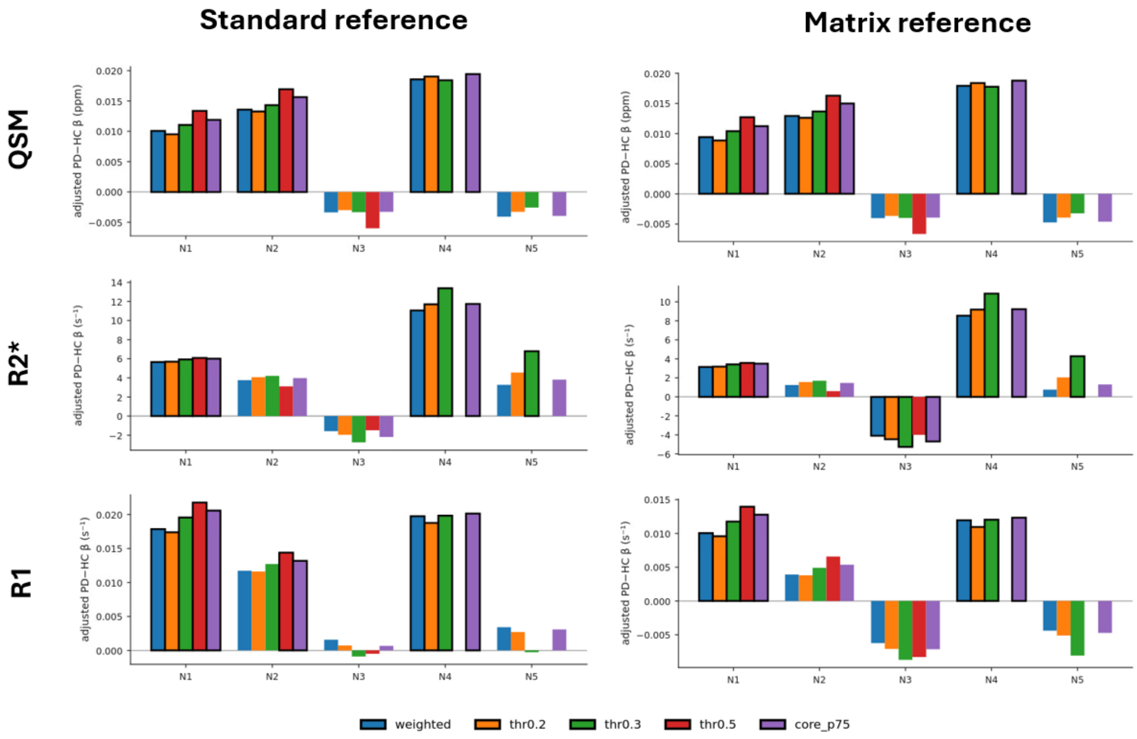


**Supplementary Fig. 3 |** Sweep of per-nigrosome effect across mask thresholds and modalities and reference schemes. Age- and sex-adjusted PD-minus-HC difference per nigrosome (N1–N5) for each modality across mask definitions: probability-weighted (weighted), binarised at p > 0.2, 0.3 and 0.5 (thr0.2, thr0.3, thr0.5), and the 75th-percentile core (core_p75). Bars with a black edge are significant at $q < 0.05$ after BH correction across N1–N5. The rank order and sign of the effects are stable across thresholds.

**Supplementary Table 11 |** Substantia nigra matrix gradient linear mixed models (LMM): group × centred bin interaction along three anatomical axes.

| Modality | Axis | Interaction β | SE | t | *P* (bootstrap) | *P* (Satterthwaite) | q (BH) | Random effects | Slope variance |
|---|---|---|---|---|---|---|---|---|---|
| QSM | Dorsoventral | 0.00036 | 0.00048 | 0.75 | 0.454 | 0.452 | 0.680 | intercept only | — |
| QSM | Rostrocaudal | −0.00273 | 0.00063 | -4.34 | <0.0001 | $6.3 \times 10^{-5}$ | 0.0003 | intercept +slope | $2.7 \times 10^{-6}$ |
| QSM | Mediolateral | 0.00013 | 0.00056 | 0.23 | 0.819 | 0.816 | 0.819 | intercept +slope | $3.7 \times 10^{-6}$ |
| R2* | Dorsoventral | 0.787 | 0.186 | 4.22 | <0.0001 | $2.6 \times 10^{-5}$ | 0.0003 | intercept only | — |
| R2* | Rostrocaudal | −0.288 | 0.263 | -1.10 | 0.287 | 0.277 | 0.389 | intercept +slope | 0.60 |
| R2* | Mediolateral | −0.296 | 0.338 | -0.88 | 0.389 | 0.385 | 0.389 | intercept +slope | 1.35 |
| R1 | Dorsoventral | 0.00024 | 0.00036 | 0.68 | 0.503 | 0.500 | 0.754 | intercept +slope | $1.3 \times 10^{-6}$ |
| R1 | Rostrocaudal | −0.00189 | 0.00041 | -4.57 | <0.0001 | $2.9 \times 10^{-5}$ | 0.0003 | intercept +slope | $1.6 \times 10^{-6}$ |
| R1 | Mediolateral | 0.00015 | 0.00057 | 0.26 | 0.793 | 0.793 | 0.833 | intercept +slope | $3.9 \times 10^{-6}$ |

Linear mixed models of matrix signal with group (PD versus HC), centred bin, their interaction, hemisphere, age and sex as fixed effects, and a random intercept and random slope of bin per subject, reduced to a random intercept where the maximal model was singular or failed to converge (n = 56, both hemispheres). Models were fitted by restricted maximum likelihood (lme4); the primary *P* is a parametric-bootstrap likelihood-ratio test of the interaction on maximum likelihood refits (pbkrtest, 10000 simulations), with the Satterthwaite approximation (lmerTest) as a sanity check. *q* from BH correction across the three axes within each modality. Interaction β is in ppm (QSM) or $s^{-1}$ (R2*, R1) per bin; slope variance is the estimated random-slope variance of bin where included.

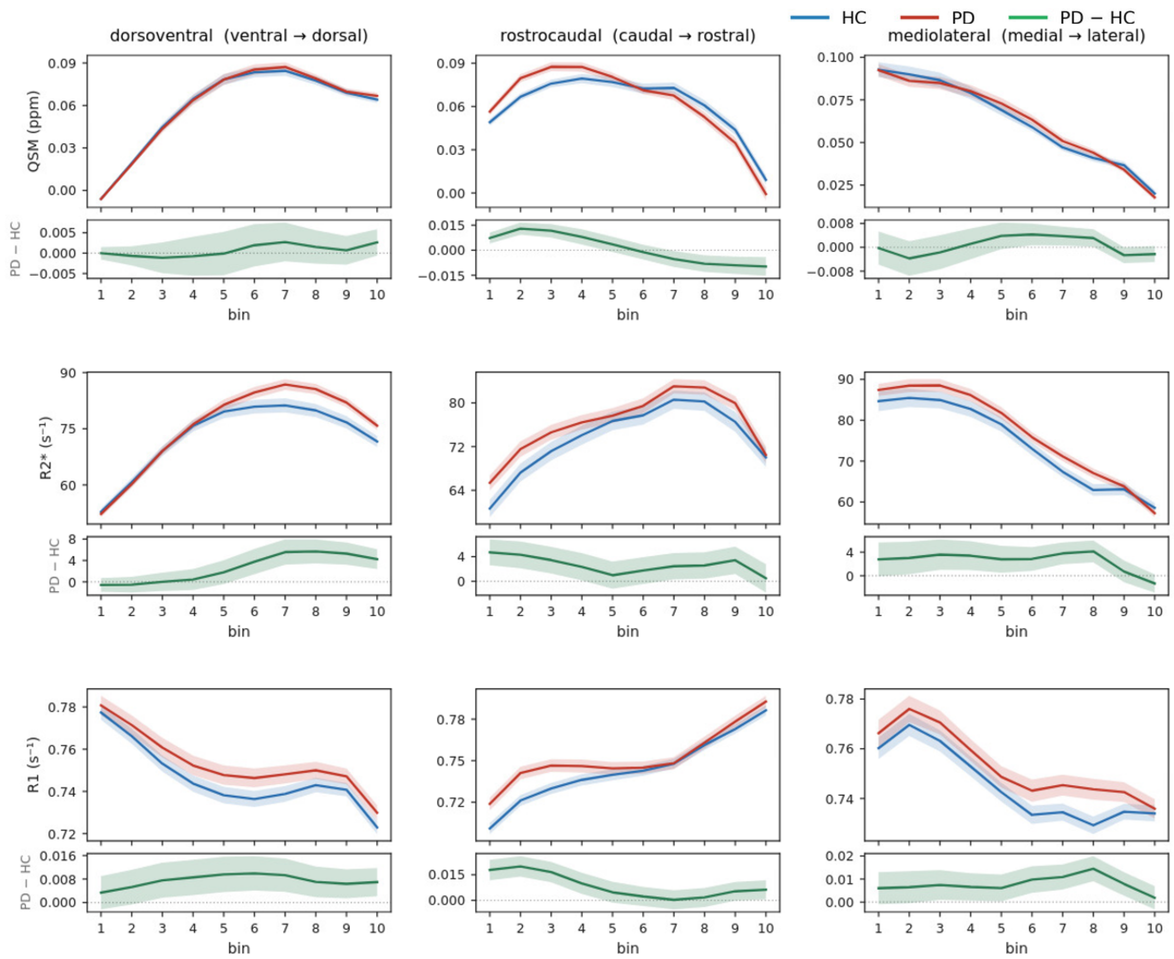


**Supplementary Fig. 4 |** Graphical summary of gradient interactions. Mean signal (with 95% confidence band) in PD (red) and HC (blue) as a function of centred bin along the dorsoventral, rostrocaudal and mediolateral axes, for QSM (top), R2* (middle) and R1 (bottom). The PD-minus-HC difference is shown in green. These profiles correspond to the interaction tests in Supplementary Table 11.

**Supplementary Table 12 |** Association between nigrosome signal and disease duration (standard reference).

| Modality | Nigrosome | Slope per year | Bootstrap 95% CI | t | Permutation P | q (BH) |
|---|---|---|---|---|---|---|
| QSM | N1 | 0.00039 | [−0.00063, 0.00138] | 0.75 | 0.441 | 0.551 |
| QSM | N2 | 0.00073 | [−0.00067, 0.00200] | 0.99 | 0.332 | 0.551 |
| QSM | N3 | 0.00070 | [−0.00003, 0.00173] | 1.56 | 0.129 | 0.440 |
| QSM | N4 | 0.00083 | [−0.00025, 0.00211] | 1.37 | 0.176 | 0.440 |
| QSM | N5 | −0.00028 | [−0.00166, 0.00061] | −0.51 | 0.610 | 0.610 |
| R2* | N1 | 0.288 | [−0.187, 1.013] | 1.10 | 0.286 | 0.428 |
| R2* | N2 | 0.137 | [−0.601, 0.600] | 0.39 | 0.701 | 0.701 |
| R2* | N3 | 0.412 | [0.013, 1.117] | 1.88 | 0.074 | 0.346 |
| R2* | N4 | 0.526 | [−0.011, 1.534] | 1.55 | 0.138 | 0.346 |
| R2* | N5 | 0.245 | [−0.142, 0.774] | 0.98 | 0.343 | 0.428 |
| R1 | N1 | 0.00060 | [−0.00082, 0.00229] | 0.88 | 0.383 | 0.806 |
| R1 | N2 | 0.00021 | [−0.00130, 0.00222] | 0.27 | 0.794 | 0.806 |
| R1 | N3 | −0.00019 | [−0.00146, 0.00148] | −0.25 | 0.806 | 0.806 |
| R1 | N4 | 0.00075 | [−0.00044, 0.00264] | 1.11 | 0.275 | 0.806 |
| R1 | N5 | 0.00042 | [−0.00055, 0.00232] | 0.66 | 0.519 | 0.806 |

Association between disease duration (years since motor-symptom onset) and bilateral nigrosome signals in the PD group (*n* = 28), adjusted for age and sex, under standard reference. Slope in ppm (QSM) or $s^{-1}$ (R2*, R1) per year. *P* from Freedman–Lane permutation; *q* from BH correction across N1–N5 within each modality.

**Supplementary Table 13 |** Association between nigrosome signal and disease duration (matrix reference).

| Modality | Nigrosome | Slope per year | Bootstrap 95% CI | t | Permutation *P* | q (BH) |
|---|---|---|---|---|---|---|
| QSM | N1 | 0.00044 | [−0.00007, 0.00075] | 2.03 | 0.054 | 0.134 |
| QSM | N2 | 0.00078 | [−0.00036, 0.00165] | 1.59 | 0.126 | 0.157 |
| QSM | N3 | 0.00075 | [−0.00015, 0.00159] | 1.60 | 0.118 | 0.157 |
| QSM | N4 | 0.00088 | [0.00019, 0.00170] | 2.35 | 0.024 | 0.118 |
| QSM | N5 | −0.00023 | [−0.00126, 0.00018] | −0.79 | 0.417 | 0.417 |
| R2* | N1 | 0.251 | [0.005, 0.670] | 1.98 | 0.062 | 0.104 |
| R2* | N2 | 0.099 | [−0.442, 0.368] | 0.48 | 0.640 | 0.640 |
| R2* | N3 | 0.374 | [0.047, 0.834] | 2.06 | 0.051 | 0.104 |
| R2* | N4 | 0.489 | [0.160, 1.226] | 2.30 | 0.031 | 0.104 |
| R2* | N5 | 0.208 | [−0.038, 0.542] | 1.28 | 0.212 | 0.266 |
| R1 | N1 | 0.00048 | [−0.00024, 0.00090] | 1.97 | 0.058 | 0.145 |
| R1 | N2 | 0.00009 | [−0.00067, 0.00095] | 0.27 | 0.794 | 0.794 |
| R1 | N3 | −0.00031 | [−0.00126, 0.00041] | −0.61 | 0.537 | 0.672 |
| R1 | N4 | 0.00063 | [0.00018, 0.00131] | 2.29 | 0.031 | 0.145 |
| R1 | N5 | 0.00030 | [−0.00031, 0.00136] | 0.77 | 0.450 | 0.672 |

Association between disease duration (years since motor-symptom onset) and bilateral nigrosome signal in the PD group (*n* = 28), adjusted for age and sex, under matrix reference. Slope in ppm (QSM) or $s^{-1}$ (R2*, R1) per year. *P* from Freedman–Lane permutation; *q* from BH correction across N1–N5 within each modality.

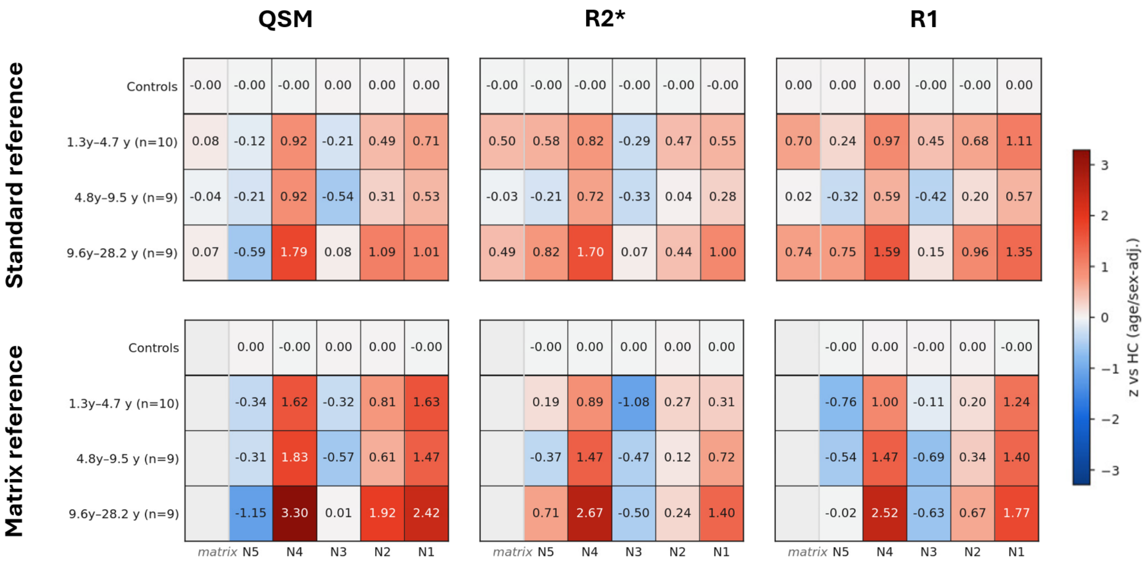


**Supplementary Fig. 5 |** Deviation from the healthy control distribution by disease-duration tertile. Heat maps of mean age- and sex-adjusted z-scores relative to the HC distribution for PD patients, divided into disease-duration tertiles, per nigrosome (N1-N5), and modality (QSM, R2*, and R1). Top: standard reference, with the matrix as an additional region. Bottom: matrix reference, with the matrix cell empty by definition. Controls are shown as a reference row (zero by construction). Colour scale is centred on zero (red, higher than HC; blue, lower than HC).

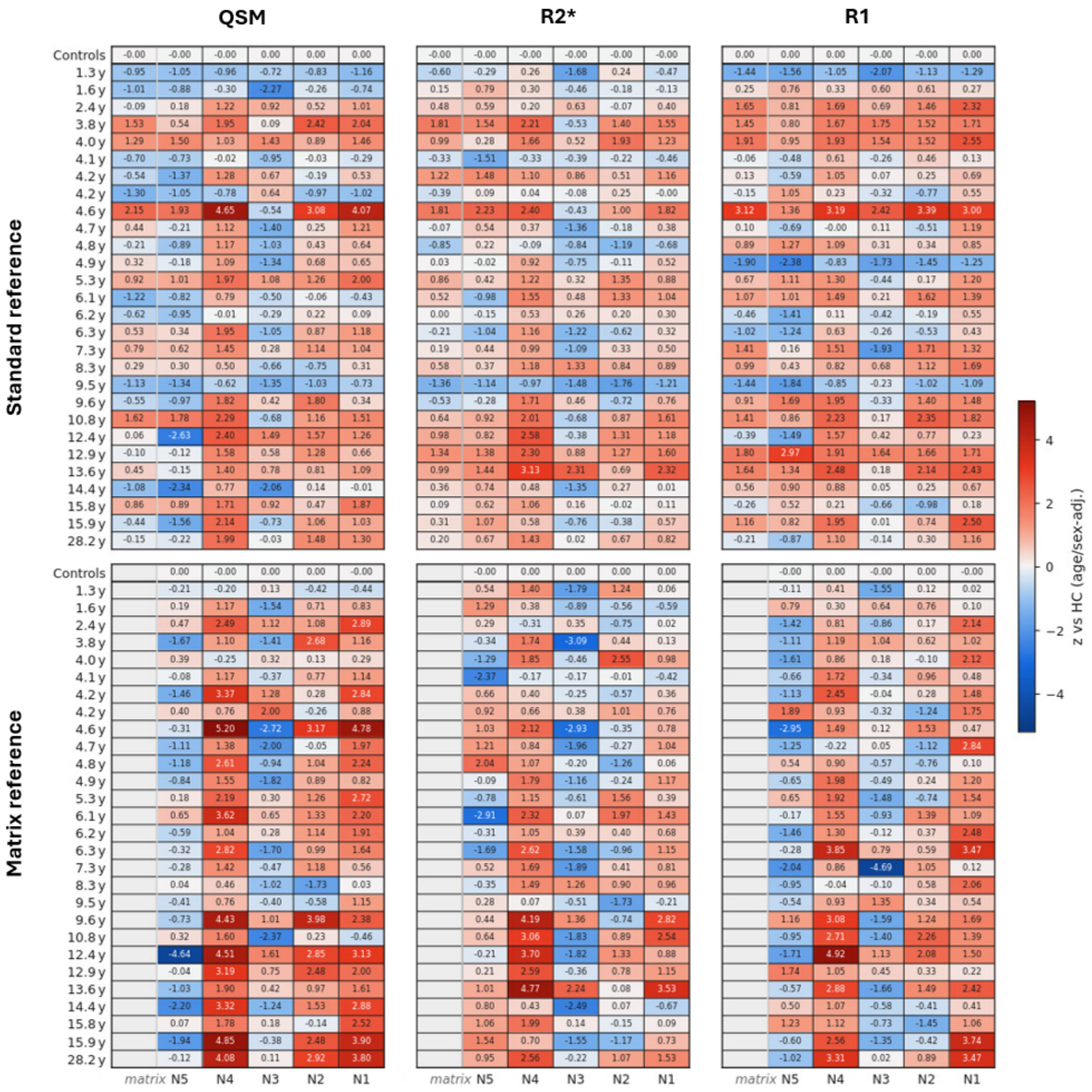


**Supplementary Fig. 6 |** Deviation from the healthy control distribution per participant, ordered by disease duration. Per-participant heatmap of age- and sex-adjusted z-scores relative to the HC distribution for PD patients, ordered by increasing disease duration (rank order; individual durations are not shown), per nigrosome (N1–N5), and modality (QSM, R2*, R1). Upper: standard reference, with the matrix as an additional region. Lower: matrix reference, with the matrix column omitted by definition. Controls are shown as a reference row (zero by construction). Colour scale centred on zero (red, higher than HC; blue, lower than HC).

**Supplementary Table 14 |** Association between imaging asymmetry and contralateral clinical motor asymmetry; primary model and leave one out stability.

| Modality | ROI | β [95% CI] | Standardized β [95% CI] | Permutation *P* | q (BH) | HC3 *P* | Jackknife sign stable | Jackknife β range | Max \|DFBETA\| |
|---|---|---|---|---|---|---|---|---|---|
| QSM | N1 | 451.8 [243.8, 733.8] | 0.645 [0.388, 0.853] | 0.0004 | 0.002 | 0.004 | Yes | [401.7, 557.3] | 105.5 |
| QSM | N2 | 59.0 [−174.7, 350.3] | 0.102 [−0.303, 0.495] | 0.653 | 0.653 | 0.683 | Yes | [1.8, 148.6] | 89.7 |
| QSM | N3 | 150.3 [−38.2, 297.3] | 0.343 [−0.063, 0.671] | 0.086 | 0.143 | 0.103 | Yes | [121.3, 187.1] | 36.7 |
| QSM | N4 | 253.0 [83.4, 392.3] | 0.538 [0.173, 0.768] | 0.004 | 0.010 | 0.003 | Yes | [212.3, 278.9] | 40.6 |
| QSM | N5 | 42.2 [−107.7, 236.0] | 0.112 [−0.294, 0.537] | 0.608 | 0.653 | 0.658 | Yes | [13.2, 111.5] | 69.3 |
| QSM | SN | 305.8 [31.8, 604.6] | 0.367 [0.031, 0.623] | 0.068 | — | 0.046 | Yes | [249.3, 387.8] | 82.1 |
| QSM | matrix | 260.2 [−9.7, 537.0] | 0.325 [−0.014, 0.596] | 0.108 | — | 0.072 | Yes | [206.8, 334.4] | 74.2 |
| R2* | N1 | 0.211 [−0.091, 0.485] | 0.316 [−0.122, 0.653] | 0.130 | 0.234 | 0.175 | Yes | [0.131, 0.282] | 0.080 |
| R2* | N2 | 0.176 [−0.074, 0.351] | 0.294 [−0.116, 0.595] | 0.140 | 0.234 | 0.104 | Yes | [0.120, 0.212] | 0.057 |
| R2* | N3 | −0.052 [−0.261, 0.236] | −0.109 [−0.593, 0.417] | 0.630 | 0.641 | 0.757 | No | [−0.114, 0.067] | 0.119 |
| R2* | N4 | 0.186 [−0.036, 0.387] | 0.369 [−0.065, 0.702] | 0.080 | 0.234 | 0.122 | Yes | [0.139, 0.252] | 0.065 |
| R2* | N5 | 0.065 [−0.217, 0.351] | 0.099 [−0.297, 0.539] | 0.641 | 0.641 | 0.647 | Yes | [0.007, 0.127] | 0.062 |
| R2* | SN | 0.300 [−0.173, 0.850] | 0.285 [−0.150, 0.679] | 0.193 | — | 0.325 | Yes | [0.181, 0.525] | 0.225 |
| R2* | matrix | 0.280 [−0.202, 0.839] | 0.263 [−0.188, 0.676] | 0.228 | — | 0.368 | Yes | [0.161, 0.510] | 0.231 |
| R1 | N1 | 38.9 [−192.0, 232.4] | 0.067 [−0.318, 0.385] | 0.766 | 0.840 | 0.714 | No | [−3.0, 71.9] | 42.0 |
| R1 | N2 | −113.1 [−311.9, 94.4] | −0.201 [−0.571, 0.171] | 0.373 | 0.840 | 0.312 | Yes | [−169.4, −75.6] | 56.3 |
| R1 | N3 | 13.9 [−129.8, 82.2] | 0.071 [−0.512, 0.431] | 0.769 | 0.840 | 0.780 | No | [−20.4, 31.0] | 34.3 |
| R1 | N4 | 18.4 [−125.3, 162.5] | 0.044 [−0.298, 0.341] | 0.840 | 0.840 | 0.793 | No | [−14.6, 36.7] | 33.0 |
| R1 | N5 | −118.7 [−321.6, 37.6] | −0.285 [−0.624, 0.088] | 0.174 | 0.840 | 0.136 | Yes | [−151.1, −92.9] | 32.4 |
| R1 | SN | −71.7 [−421.9, 215.6] | −0.085 [−0.425, 0.254] | 0.688 | — | 0.678 | No | [−146.1, 30.8] | 102.5 |
| R1 | matrix | −78.3 [−446.9, 209.2] | −0.092 [−0.438, 0.251] | 0.664 | — | 0.657 | No | [−155.0, 31.8] | 110.2 |

Regression of the lateralised MDS-UPDRS part III asymmetry index on the contralateral imaging asymmetry of each region in the PD group ($n = 28$), adjusted for age and sex. β is the change in the clinical asymmetry index per unit imaging asymmetry (ppm for QSM; $s^{-1}$ for R2*, R1), with bootstrap 95% CI; standardised β is in SD units. *P* from Freedman–Lane permutation; *q* from BH correction across N1–N5 within each modality (SN and matrix shown for context); HC3 *P* uses heteroscedasticity-consistent standard errors. Jackknife columns report whether the sign of β was preserved in all leave-one-out refits, the range of the leave-one-out β estimates, and the largest absolute change in β when a single participant was omitted (DFBETA). Dashes in the q column denote rows excluded from the correction family.

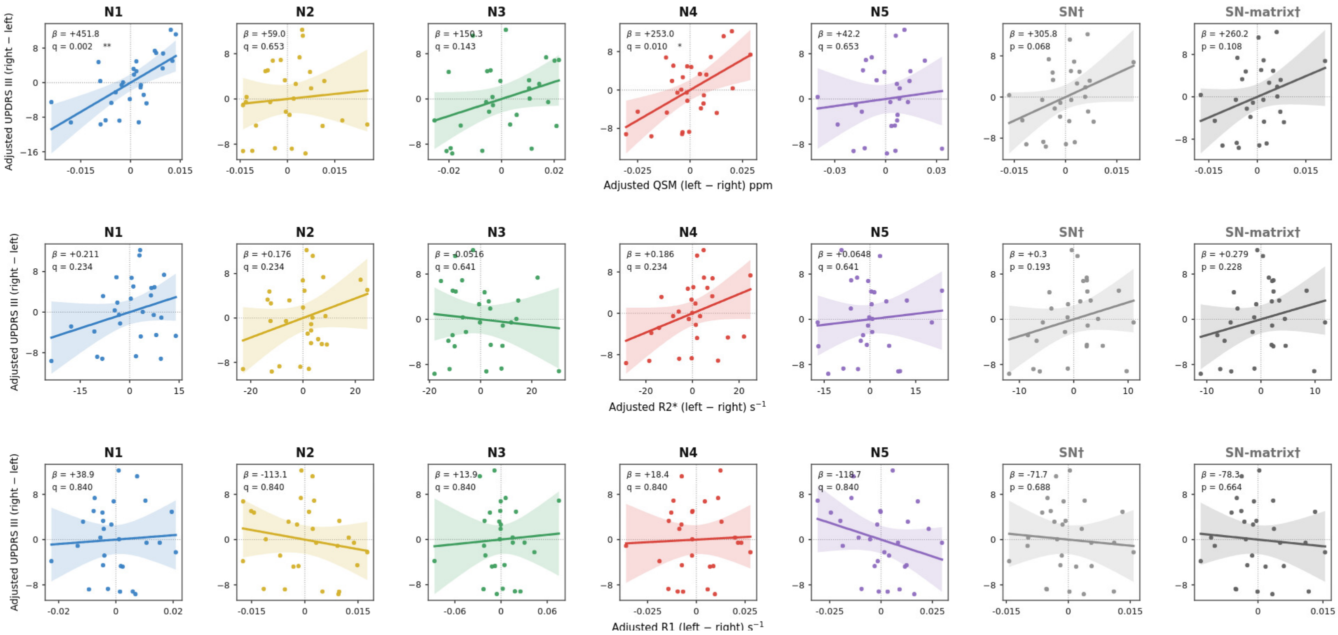


**Supplementary Fig. 7 |** Motor asymmetry summary plots. Partial regression plots of the clinical asymmetry index against contralateral imaging asymmetry for each region, after adjustment for age and sex (PD group, $n$ = 28), for QSM (top), R2* (middle) and R1 (bottom). Lines show the fitted slope with 95% confidence band. Numerical results are given in Supplementary Table 14.

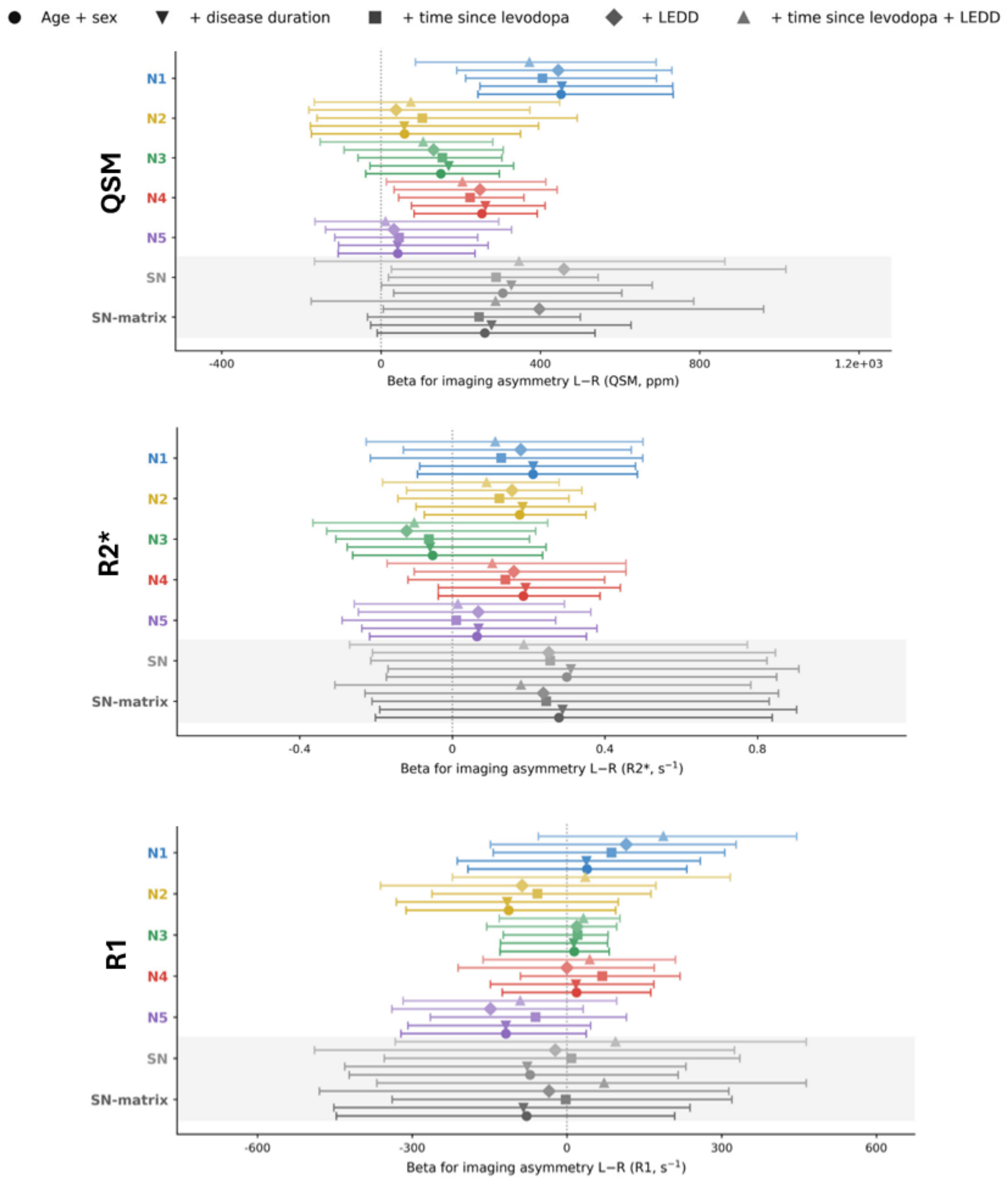


**Supplementary Fig. 8 |** Motor asymmetry, covariate sensitivity. β estimates with bootstrap 95% CI for each region under the primary model (age + sex) and four sensitivity models (disease duration, time since levodopa, levodopa equivalent daily dose (LEDD), and time since levodopa + LEDD) for QSM, R2*, and R1. Marker shape denotes the covariate set.
